# Widespread post-transcriptional regulation of co-transmission

**DOI:** 10.1101/2023.03.01.530653

**Authors:** Nannan Chen, Yunpeng Zhang, Emmanuel J. Rivera-Rodriguez, Albert D. Yu, Michael Hobin, Michael Rosbash, Leslie C. Griffith

## Abstract

While neurotransmitter identity was once considered singular and immutable for mature neurons, it is now appreciated that one neuron can release multiple neuroactive substances (co-transmission) whose identities can even change over time. To explore the mechanisms that tune the suite of transmitters a neuron releases, we developed transcriptional and translational reporters for cholinergic, glutamatergic, and GABAergic signaling in *Drosophila*. We show that many glutamatergic and GABAergic cells also transcribe cholinergic genes, but fail to accumulate cholinergic effector proteins. Suppression of cholinergic signaling involves posttranscriptional regulation of cholinergic transcripts by the microRNA miR-190; chronic loss of miR-190 function allows expression of cholinergic machinery, reducing and fragmenting sleep. Using a “translation-trap” strategy we show that neurons in these populations have episodes of transient translation of cholinergic proteins, demonstrating that suppression of co-transmission is actively modulated. Posttranscriptional restriction of fast transmitter co-transmission provides a mechanism allowing reversible tuning of neuronal output.

**One-Sentence Summary:** Cholinergic co-transmission in large populations of glutamatergic and GABAergic neurons in the *Drosophila* adult brain is controlled by miR-190.

---

Small molecule chemicals mediating neuronal communication are packaged into vesicles for release by vesicular neurotransmitter transporter proteins (vNTs). The most common fast-acting neurotransmitters in both vertebrates and invertebrates each have a cognate vNT (or vNT family): VAChT for acetylcholine (ACh), VGAT for gamma-amino butyric acid (GABA) and VGluT for glutamate (Glu) (*1*). Co-transmission, release of multiple neuroactive molecules from a single cell, has been reported in many animals, and usually involves release of a bioamine or peptide neuromodulator with a fast transmitter (*2, 3*). This type of modulation can be regulated by changes in environment or neuronal activity (*4*). Interestingly, co-transmission between multiple fast-acting neurotransmitters has only been seen functionally in a few cases (*5, 6*), though some studies have reported the co-expression of multiple vNT mRNAs (*7–9*). Such co-transmission can have profound effects on circuit dynamics (*10, 11*). Using new genetic tools to study transcription and translation of vNTs for fast neurotransmitters, we demonstrate here that there are large populations of fully differentiated glutamatergic and GABAergic neurons in the adult fly brain that transcribe genes specifying synthesis and release of ACh but block accumulation of protein products via microRNA (miR) repression. This suggests a widespread but tightly-regulated potential for co-transmission.

To map the extent of co-transcription of vNTs, we used a split-Gal4 strategy (*12*) in which Gal4-DBD or AD sequences are inserted into the endogenous loci of *VAChT*, *VGluT* and *VGAT* genes to put them under control of NT-specific transcriptional programs (Fig. 1A and fig. S1A). Both the *VGluT-AD:VAChT-DBD* (Fig. 1B) and *VAChT-AD:VGluT-DBD* (fig. S1C) split-Gal4s show broad expression with the strongest signal in fan-shape body (FSB) neurons. As expected for intersectional drivers, *VAChT:VGluT* split-Gal4 labels fewer neurons than either *VAChT-* or *VGluT-Gal4* drivers (fig. S1B). We will refer to the cell subset labeled by this intersectional tool as “Glu^ACh^” neurons and the split-Gal4 as *Glu^ACh^-Gal4*. Similarly, both the *VGAT-AD:VAChT-DBD* (Fig. 1C) and *VAChT-AD:VGAT-DBD* (fig. S1D) split-Gal4s had a broad, but distinct expression profile with the strongest EGFP signal in ellipsoid body (EB) neurons; we call these cells “GABA^ACh^” neurons. *VGAT:VGluT* split-GAL4 brains showed little consistent co-expression (data not shown). These results suggested potential co-expression of *VAChT* with both the *VGluT* and *VGAT* genes and possible co-transmission.

**Fig. 1.**
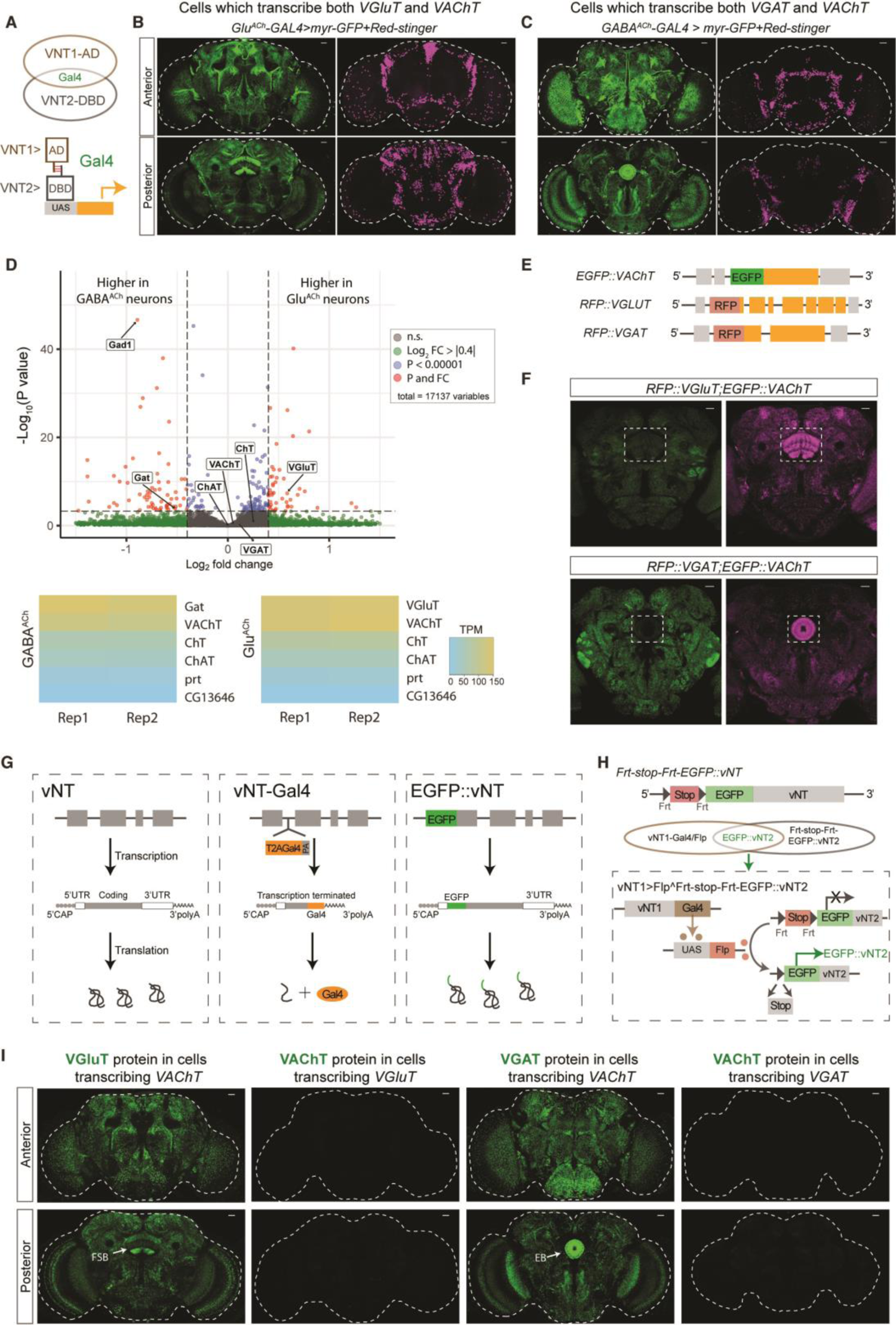
Transcription of VAChT in VGluT/VGAT positive neurons. (**A**) Schematic diagram of split-Gal4 strategy. Expression of AD and DBD in the same neurons reconstitutes Gal4 protein to initiate expression. (**B-C**) Expression patterns of *VGluT-AD;VAChT-DBD split-Gal4* and (**B**) *VGAT-AD;VAChT-DBD split-Gal4* (**C**): Anterior (top) and posterior (bottom). Green indicates neuronal membrane, while magenta shows nuclei. Dashed white lines outline the brain. (**D**) Nuclear RNAseq demonstrates high cholinergic mRNA levels in Glu^ACh^ and GABA^Ach^ neurons. Volcano plot (top) shows statistically significant enrichment of *VGluT* in Glu^ACh^ cells (adjusted P value (P_adj_) < 0.05; log_2_ fold change (FC) = 0.58) and Gat (P_adj_ < 0.05; log_2_FC = −0.52) and Gad1 (P_adj_ < 0.05; log_2_FC = −0.89) in GABA^ACh^ neurons. *VAChT, ChaT* and *ChT* mRNAs were not differentially expressed (P_adj_ > 0.05). Heat maps (bottom) for each cell type show that cholinergic markers are present in both cell types at levels (TPM, transcripts per million) comparable to *VGluT* and *VGAT* respectively, while control vesicular transporters are not expressed. (**E**) Schematic diagram of N-terminal genomic fusion lines *EGFP::VAChT, RFP::VGluT* and *RFP::VGAT.* (**F**) Representative single-slice pictures of adult brains of *RFP::VGluT;EGFP::VAChT* (left) and *RFP::VGAT; EGFP::VAChT* (right) flies. Green indicates EGFP expression while magenta indicates RFP expression. The dashed box outlines EGFP and RFP signals in fan-shape body (left) and ellipsoid body (right). (**G**) Schematic diagrams showing the transcription and translation processing of vNT mRNA in wildtype (left), T2A Gal4 alleles (middle) and EGFP::vNT fusion alleles (right). For vNT-Gal4 alleles, transcription of vNT is terminated at the Gal4 insertion site, inducing loss of vNT 3’UTR information, with production of separate terminated vNT and GAL4 proteins. For EGFP::vNT alleles, EGFP is transcribed and translated within the intact vNT mRNA, making EGFP::vNT fusion protein. (**H**) Schematic diagram showing the flip-out stop gene strategy. vNT1-Gal4 drives FLP recombinase expression which excises the FRT-flanked stop cassette preceding EGFP::vNT2, allowing EGFP::vNT2 expression. Thus, the EGFP signals indicate transcription of vNT1 with transcription and translation of vNT2 in the same neuron. (**I**) *VAChT-Gal4* flip-out derepression of EGFP::VGluT shows EGFP in FSB, while *VAChT-Gal4* flip-out of EGFP::VGAT shows EGFP in EB. Flip-out derepression ECFP::VAChT shows no signal. Dashed white lines indicate the whole brain. Scale bars = 20µm.

To verify co-transcription of the native vNT genes in these cells, we analyzed nuclear polyA-containing RNA from INTACT-sorted Glu^ACh^ and GABA^ACh^ nuclei (*13*) (Fig. 1D), a technique which minimizes the effects of cytoplasmic posttranscriptional processes on mRNA levels (*14*). GABA^ACh^ nuclei express high levels of *GAD1* and *GAT* mRNA, while Glu^ACh^ nuclei express high levels of *VGluT* as expected*. VAChT*, *ChaT and ChT* mRNA are also expressed strongly in both cell types. Surprisingly, nuclear *VGAT* mRNA was also found in both cell types. *Portabella* and *CG13646*, vNTs related respectively to VAChT/VGluT and VGAT, were not found at significant levels in either population.

To be co-transmitting, Glu^ACh^ and GABA^ACh^ neurons would need to express the protein products of both vNT genes. To directly visualize the vNT proteins we fused fluorescent proteins (FPs) to the N-termini of the endogenous coding sequences using CRISPR/Cas9 (Fig. 1E). These fusion alleles faithfully recapitulate the native protein distribution as assessed by immunostaining of heterozygotes (fig. S2). Co-staining for EGFP and RFP in *RFP::VGluT;EGFP::VAChT* fly brains, we found strong RFP::VGluT protein expression in FSB neurons, but no EGFP::VAChT protein at the same level of the confocal stack (Fig. 1F). Similarly, in *RFP::VGAT;EGFP::VAChT* fly brains, strong RFP::VGAT staining is present in EB neurons, but EGFP::VAChT protein is not (Fig. 1F).

While split-Gal4 expressed from the *VAChT* locus is clearly present in FSB and EB, the lack of EGFP::VAChT indicates that the protein does not accumulate in these regions. We hypothesized that difference may be a function of the structure of the *VAChT* transcripts produced in these two different CRISPR-engineered animals. In split-GAL4 lines (and the T2A-Gal4 lines used below), GAL4 coding sequence(s), followed by a polyadenylation site, are inserted into a vNT intron, producing a truncated transcript that lacks the vNT gene’s 3’UTR, a region which can contain *cis* regulatory sequences regulating translation and/or RNA stability (*15*). For FP::vNT fusion alleles, the FP coding sequence is fused in-frame to form a functional chimeric vNT protein, meaning the FP::vNT mRNA has all the regulatory information native to the wildtype vNT mRNA (Fig. 1G). This suggests that while both the *VAChT* split-Gal4 and EGFP::VAChT mRNA are transcribed in Glu^ACh^ and GABA^ACh^ neurons but that mRNA containing native 3’UTR sequences is not translated.

To test this idea, we created conditional FP fusion alleles containing an *Frt-stop-Frt-FP* cassette downstream of the start codon of each vNT gene (Fig. 1H). In these animals, FP::vNT transcription is blocked until FLP recombinase is expressed, excising the stop cassette. GAL4+ cells then become competent to generate a FP::vNT mRNA containing all the endogenous UTR information. *Frt-stop-Frt-ECFP::VAChT* flies were validated by driving FLP expression with *VT030559-Gal4* in cholinergic mushroom body cells. ECFP::VAChT was present in mushroom body as expected and dependent on GAL4 (fig. S3A). Similarly, EGFP::VGluT in FSB neurons and EGFP::VGAT signals in the anterior paired lateral (APL) neurons demonstrate the specificity of these lines (fig. S3B and C).

Fig. 1H shows the strategy used to test for posttranscriptional suppression of VAChT protein expression in Glu^ACh^ and GABA^ACh^ neurons. FLP recombinase, driven in cells which transcribe vNT1, catalyzes excision of the stop cassette for FP::vNT2. Only if the cells which transcribe vNT1 are also competent to both transcribe and translate vNT2, is an FP signal is seen. Using *VAChT-Gal4* to flip out the stop cassette for *EGFP::VGluT* results in strong protein signal in the same pattern observed for *Glu^ACh^-Gal4*, indicating that *VGluT* is both transcribed and translated in this subset of *VAChT*-transcribing neurons (Fig. 1I). However, FLP-derepression of *ECFP:: VAChT* with *VGluT-Gal4* produces no detectable protein in adult brain (Fig. 1I), suggesting either degradation or translational suppression of *VAChT* mRNA in Glu^ACh^ cells. GABA^ACh^ neurons behaved similarly: *EGFP::VGAT* expression confirmed transcription and translation of *VGAT* mRNA while the absence of ECFP::VAChT protein shows there is no translation of *VAChT* mRNA in GABA^ACh^ neurons. Thus, *VGluT/VGAT* are transcribed and translated in Glu^ACh^/ GABA^ACh^ neurons while *VAChT* mRNA is transcribed, but either degraded or untranslated, in both groups.

Control of protein synthesis by microRNAs, small non-coding RNAs, which bind to mRNA to initiate degradation or inhibit translation, is widespread (*16*). *In silico* evaluation of the *VAChT* 3’UTR (www.targetscan.org/fly_72/) identified multiple high-confidence binding sites for miR-190, a microRNA which also targets several other cholinergic mRNAs, including *ChAT,* acetylcholine’s synthetic enzyme and *ChT,* the choline transporter. To determine whether miR-190 regulates production of these cholinergic effector proteins, we assayed its ability to suppress expression of a Firefly luciferase (Fluc) gene which had either the *VAChT* or *ChAT* 3’UTR. Co-transfection of S2 cells with miR-190 and a *Renilla* luciferase (Rluc) control plasmid induces a significant decrease in the Fluc/Rluc ratio with both 3’UTRs compared to a scrambled miR, but no miR-190-dependent decrease is found when the three putative miR-190 binding sites of the *ChAT* 3’UTR are deleted or there is no 3’UTR (Fig. 2A). These results suggest that miR-190 can suppress the expression of cholinergic proteins by directly binding to their 3’UTRs.

**Fig. 2.**
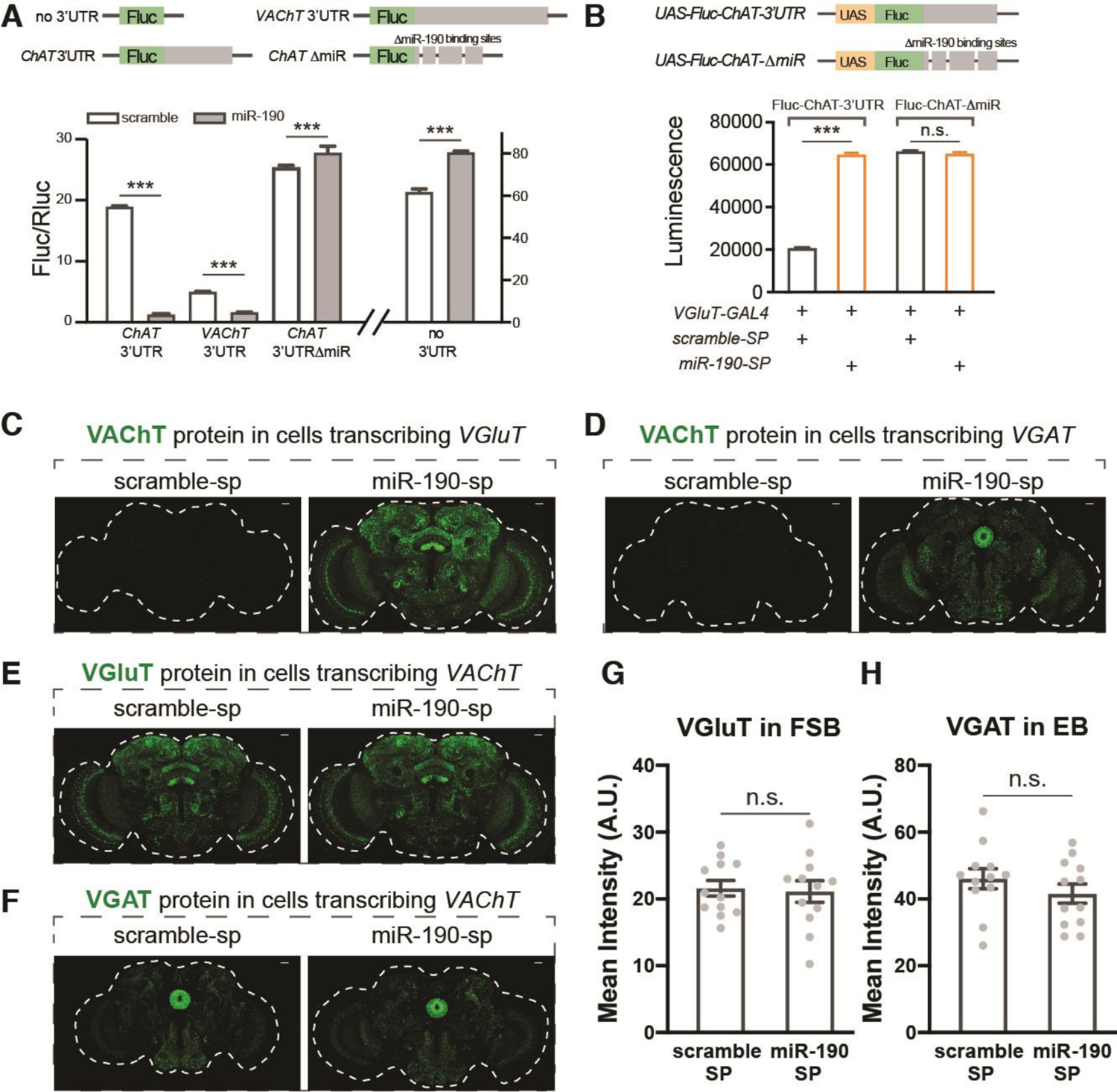
MiR-190 blocks VAChT protein accumulation. (**A**) S2 cells were co-transfected with Fluc-UTR, Rluc and scramble or miR-190 plasmids. For the no 3’UTR plasmid, only Fluc is included; for the VAChT-3’UTR and ChAT-3’UTR plasmids, Fluc is followed by 3’UTR of *VAChT* or *ChAT*; for the ChAT-del plasmid, the three predicted miR-190 binding sites in the *ChAT* 3’UTR are deleted. Firefly luciferase (Fluc) activity was normalized to *Renilla* luciferase (Rluc) activity. n=6 for each group. Co-expression of miR-190 blocks Fluc expression only when plasmids contain *VAChT* or *ChAT* 3’UTRs. Deletion of predicted miR-190 binding sites in the *ChAT* 3’UTR blocks miR-190 suppression. (**B**) Expression of miR-190 sponge in VGluT positive neurons up-regulates luciferase activity when the Fluc transgene has a *ChAT* 3’UTR, indicating that miR-190 is expressed in these adult neurons. When miR-190 binding sites are deleted from the transgene’s *ChAT* 3’UTR, luciferase activity is no longer responsive to miR-190 sponge. n=6 for each group. (**C-D**) Representative pictures of *VGluT-Gal4* (**C**) or *VGAT-Gal4* (**D**) driving flip-out derepression of *ECFP::VAChT* flies, with scramble or miR-190 sponge expressed in the same neurons. (**E-F**) Representative pictures of *VAChT-Gal4* driving flip-out derepression of *EGFP::VGluT* (**E**) or *EGFP::VGAT* (**F**) while expressing scramble or miR-190 sponge in the same neurons. (**G**) Quantification of EGFP::VGluT protein in FSB neurons from panel **E**. (**H**) Quantification of EGFP::VGAT protein in EB neurons from panel **F**. n=12 for each group in panels **G** and **H**. Dashed white lines indicate the whole brain. Scale bars = 20µm. Data are shown as mean ± SEM, and analyzed by Student’s t-test. n.s. indicated no difference, *** indicates p<0.001. Gray dots show individual values in panels **G** and **H**.

To test the idea that miR-190 is responsible for *in vivo* suppression of cholinergic transmission in Glu^ACh^ and GABA^ACh^ cells, we suppressed miR-190 function in specific neurons using miR-190 sponge lines (*17*). To validate the specificity and efficacy of the sponges and to test for the presence of miR-190 in adult glutamatergic cells, we created UAS-driven Fluc reporter lines that had either the *ChAT* 3’UTR or a mutant *ChAT* 3’UTR with miR-190 sites deleted. Co-expression of the reporters under control of *VGluT-Gal4* with the miR-190 sponge or scramble control demonstrates that miR-190 is present in adult glutamatergic neurons and that its function can be inhibited *in vivo* by the sponge (Fig. 2B).

To explore the role of miR-190 in regulation of endogenous VAChT translation, we asked if expression of the miR-190 sponge would result in ECFP::VAChT protein expression in glutamatergic or GABAergic neurons. For both Glu^ACh^ and GABA^ACh^ cells, miR-190 sponge produced strong ECFP::VAChT protein signal in the expected patterns (Fig. 2C and D and fig. S4). These results indicate that miR-190 suppresses accumulation of *VAChT* protein in Glu^ACh^ and GABA^ACh^ cells in adult heads. Interestingly, expression of miR-190 sponge does not change the expression pattern or intensity of EGFP::VGluT or EGFP::VGAT protein in FSB or EB neurons (Fig. 2E to H), suggesting that miR-190 has no role in the regulation of VGluT or VGAT protein levels.

The circuitry controlling sleep in *Drosophila* includes regions that contain Glu^ACh^ (dFSB) and GABA^ACh^ (EB) neurons (*18*). Suppression of miR-190 function pan-neuronally, as well as in either glutamatergic or cholinergic neurons, reduces daytime and nighttime sleep significantly (fig. S5A to C) compared to expression of a scrambled control sponge. To ask if this is the result of reducing miR-190 in Glu^ACh^ neurons, we expressed the sponge under control of *ChAT-GAL4* with *VGluT-GAL80* to block GAL4 action in Glu^ACh^ cells, and found the sleep reduction was rescued (Fig. 3A). Indeed, suppression of miR-190 function in Glu^ACh^ neurons using Glu^ACh^-GAL4 split drivers also leads to a large reduction in total sleep (Fig. 3B). Taken together, this demonstrates that loss of miR-190 in Glu^ACh^ cells decreases sleep. Suppression of miR-190 function in GABAergic neurons (fig. S5D) or specifically in GABA^ACh^ cells (Fig. 3C) also decreases nighttime sleep, but to a lesser extent than Glu^ACh^ manipulations. However, there is significant sleep fragmentation with miR-190 sponge in both populations (Fig. 3A to C and fig. S6 and fig. S7). Locomotor activity while awake is unaffected or reduced (fig. S8), indicating the decrease of sleep is not due to hyperactivity.

**Fig. 3.**
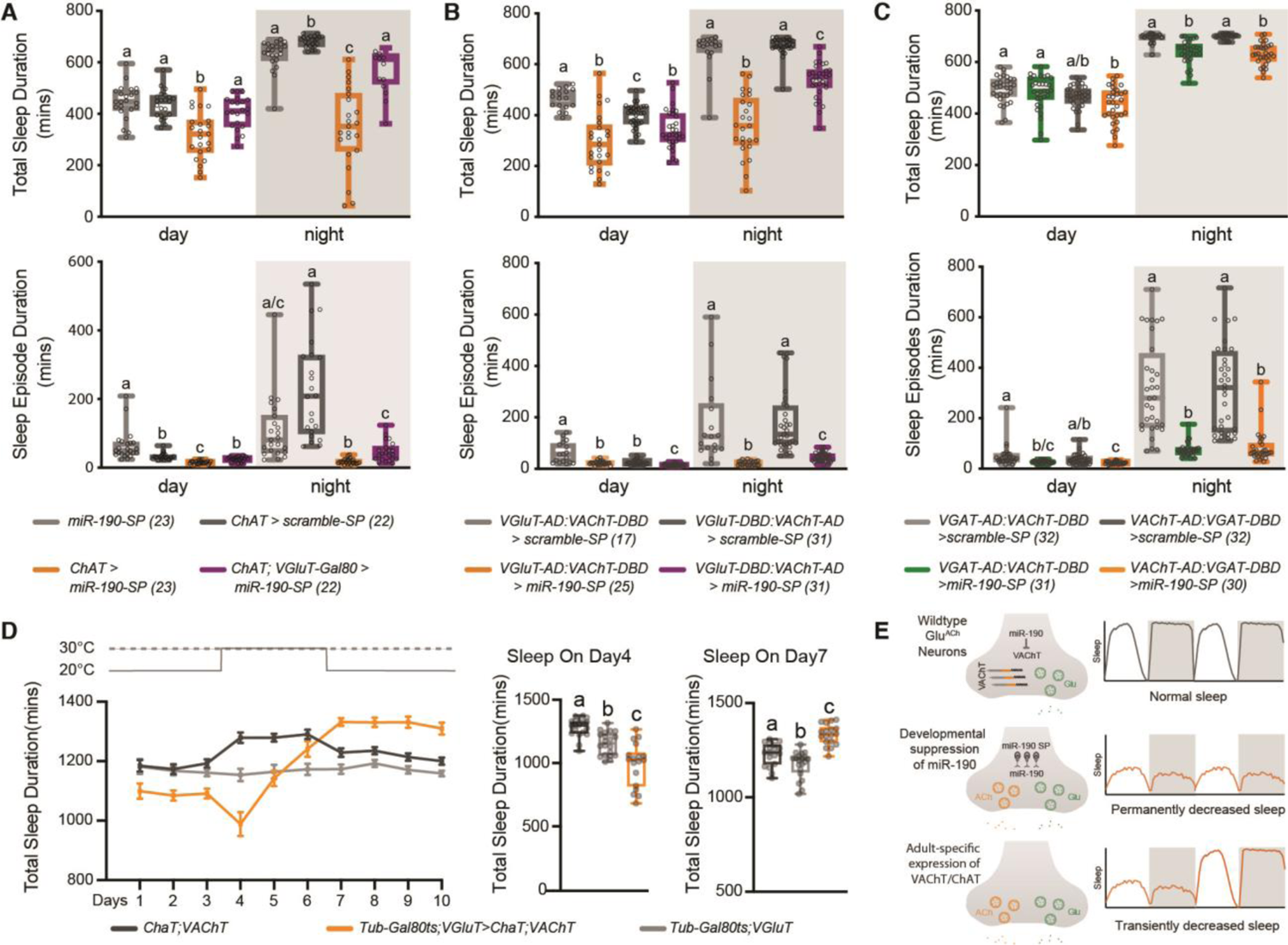
MiR-190 regulates sleep by controlling cholinergic co-transmission in glutamatergic neurons. (**A**) Reduction and fragmentation of sleep by suppression of miR-190 function in cholinergic neurons maps to neurons also expressing VGluT. (**B**) In *VGluT:VAChT* split-Gal4 neurons, miR-190 suppression reduces and fragments daytime and nighttime sleep. (**C**) In *VGAT:VAChT* split-Gal4 neurons, miR-190 suppression reduces and fragments daytime and nighttime sleep. (**D**) Temporally-controlled overexpression of ChAT and VAChT from transgenes lacking cognate 3’UTRs in VGluT+ neurons decreases sleep acutely and triggers fast compensation. n=18-20. Data are shown as mean ± SEM, gray circles show individual values. Statistical differences are indicated by letters, with genotypes that are not significantly different having the same letter. (**E**) Summary model. In Glu^ACh^ neurons, suppression of miR-190 function during development induces ACh co-transmission and alters adult sleep circuits. Adult-specific expression of VAChT/ChAT decreases sleep acutely and triggers strong homeostatic compensation.

We reasoned that if the sleep effects of miR-190 suppression were due to cholinergic transmission in glutamatergic neurons, expressing both ChAT and VAChT in these neurons should phenocopy the miR-190 sponge. Although *Glu^ACh^-GAL4*-driven expression of ChAT/VAChT transgenes lacking 3’UTR sequences was completely lethal (supporting the importance of the miR-190 suppression mechanism), limiting expression to adulthood with *TubGAL80^ts^* rescued viability and was sufficient to immediately both decrease and severely fragment sleep (Fig. 3D and fig. S9A and B). But in contrast to the suppression of miR-190 function using drivers that express during development, total sleep recovers rapidly, even before the end of protein induction (Fig. 3D and fig. S9C). These data suggest that the adult sleep phenotype seen with temporally-uncontrolled expression of miR-190 sponge may be due to developmental rewiring of the sleep homeostat circuit; limiting sponge expression using *Tub-GAL80^ts^* supports this (Rivera-Rodriguez and Adel et al., in preparation). We hypothesize that without developmental suppression of miR-190 function, the homeostat is intact and the sleep loss due to adult expression of VAChT/ChAT is subject to strong compensation (Fig. 3E).

The adult persistence of the miR-190 mechanism for suppression of cholinergic transmission raises the question of whether there are situations where co-transmission is permitted as a form of plasticity. While VAChT accumulation in normal Glu^ACh^ or GABA^ACh^ cells is not detectable (Fig. 1I), limited local or transient expression would be difficult to visualize. To capture transient events, we designed a “translation-trap” (Fig. 4A). The FLP recombinase coding region was inserted into the *VAChT* locus. FLP-encoding mRNA is translated only under conditions permissive for *VAChT* mRNA translation. Combining this allele with *FRT-stop-FRT-EGFP::VGluT* and *VGAT* alleles allows permanent marking of Glu^ACh^ and GABA^ACh^ neurons that have at some point translated *VAChT* mRNA. Fig. 4B shows EGFP::VGluT staining indicating there are Glu^ACh^ neurons in the pars intercerebralis and ventral areas of the brain which have translated *VAChT* mRNA. Similarly, EGFP::VGAT signals in EB, medulla and several other central brain regions demonstrate translation of *VChAT* in GABA^ACh^ neurons (Fig. 4C). These data show that miR-190 function is transiently suppressed in multiple neuron groups.

**Fig. 4.**
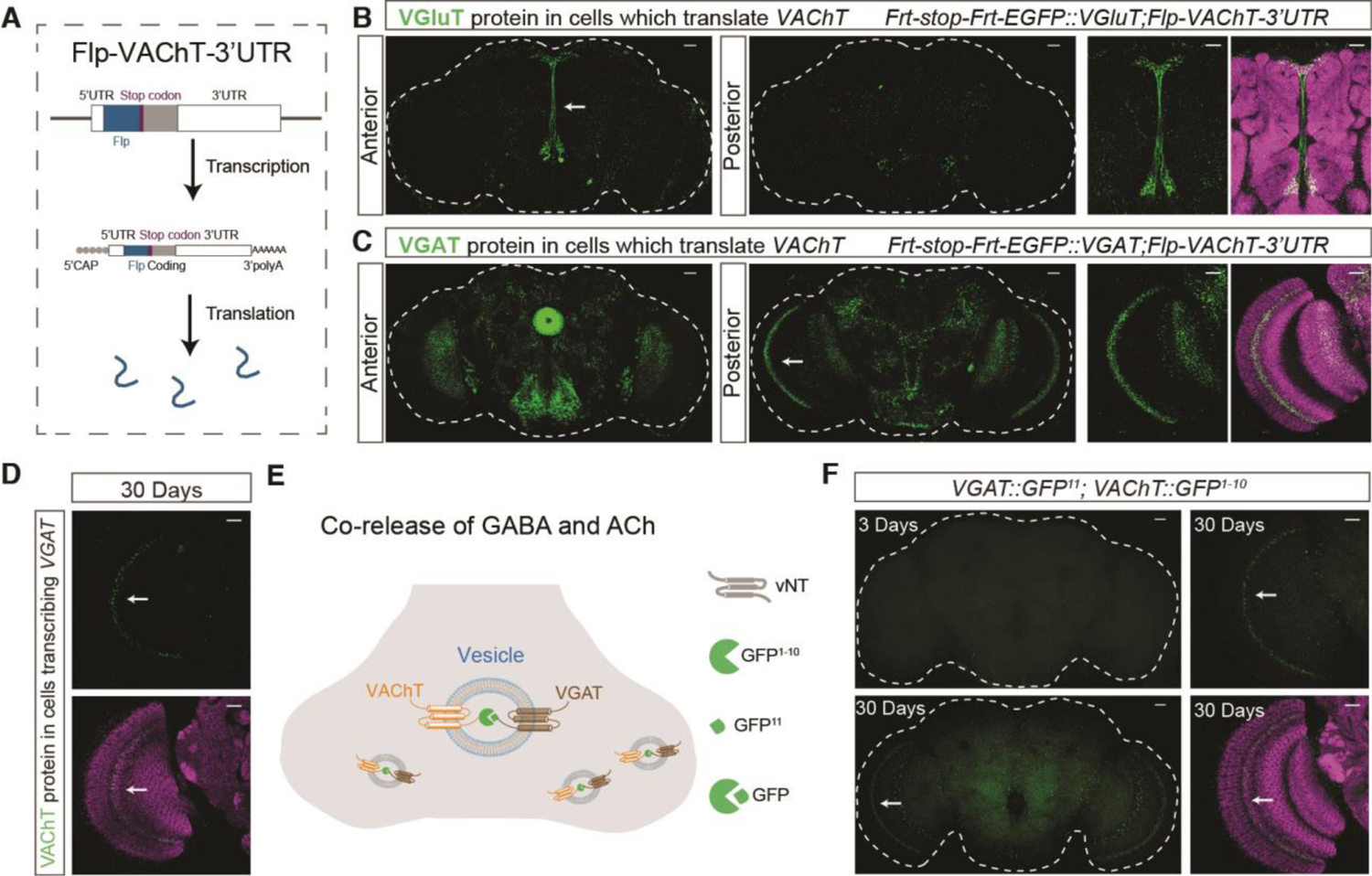
VAChT repression is released in specific cell types and VAChT traffics to VGAT vesicles. (**A**) Schematic diagram showing the translation-trap strategy. In *Flp-VAChT-3’UTR* flies. *Flp* is transcribed and translated as part of the full 3’UTR-containing *VAChT* mRNA. (**B**) *Flp-VAChT-3’UTR* flip-out derepression of *EGFP::VGluT* marks central brain neurons. White arrow shows the region enlarged at right. (**C**) *Flp-VAChT-3’UTR* flip-out derepression of *EGFP::VGAT* medulla and central brain neurons. White arrow shows the region enlarge at right. (**D**) In 30 day old flies, *VGAT-Gal4* flip-out derepression of *ECFP::VAChT* (strategy as in Fig. 1H) generates ECFP signal in medulla (white arrow). (**E**) Schematic of strategy to visualize VAChT localization. *VAChT* and *VGAT* alleles were generated with lumenal split GFP fusions. GFP reconstitution only occurs if VAChT and VGAT are in the same vesicle. (**F**) In 30 day old flies, reconstituted GFP signal is visible without staining in medulla neurons. In 3 day old flies, no GFP is detected. For panels **B** to **D and panel F**, green shows EGFP or ECFP, while magenta is Brp staining. Scale bars = 20µm.

Because our translation-trap is an irreversible mark, it does not indicate when or for how long VAChT translation occurred, or whether it is responsive to physiological state. To ask whether VAChT translation was occurring in adults, we returned to animals in which VAChT in GABA^ACh^ cells is tagged with ECFP (Fig. 1H). While ECFP::VAChT was undetectable in young animals (Fig. 1I), it appears in GABA^ACh^ medulla neurons in 30-day old brains, consistent with results from the translation-trap showing these cells translate VAChT (Fig. 4D). VGAT expression in GABA^ACh^ neurons did not change with age (fig. S10). These data demonstrate that VAChT translation occurs in mature GABA^ACh^ neurons and is stimulated by physiological changes associated with aging.

Though the appearance of VAChT protein in nerve terminals is consistent with ability to package ACh, we sought to determine if the protein was in synaptic vesicles. We knocked *GFP^1-10^* and *GFP^11^* into the *VAChT* gene, and *GFP^11^* into the *VGAT* gene (fig. S11A and B), such that the split GFP would be on the luminal face of the vNT (fig. S11C). Notably, in 30-day-old flies, we found clear reconstitution of live GFP signals between VAChT-GFP^1-10^ with VGAT-GFP^11^ in medulla neurons; no signal was found 3-day-old animals (Fig. 4E and F). These results indicate that VAChT and VGAT are present in the same vesicles, suggesting ACh and GABA co-release in aging flies. While the functional effect of this co-release has yet to be determined, it is notable that aging in flies, like in humans, is associated with significant increases in sleep fragmentation (*19*).

Co-transmission is now recognized as a common and important mode of neuronal communication, and it can be dynamic. NT plasticity involving replacement of one transmitter with another, either developmentally (*20*) or in the context of a few neurons in a mature circuit (*4, 21*), has been shown in multiple species. In cases where the molecular mechanism is known, these switching events have ultimately required transcriptional changes (*22, 23*). In this study we describe a mechanistically-distinct phenomenon in which the transcription of cholinergic genes is already active in thousands of GABAergic and glutamatergic neurons in the adult fly brain, and functional expression is controlled by a reversable microRNA switch. Since GABA and glutamate are generally inhibitory transmitters in the central brain of *Drosophila,* this gives Glu^ACh^ and GABA^ACh^ neurons the ability to rapidly and transiently alter the magnitude or even the sign of their output by scaling miR-190 levels. These neurons may be akin to the reserve pool neurons of the adult zebrafish spinal cord that can reversibly acquire and release glutamate to enhance neuromuscular junction function acutely after locomotor stress (*24*).

While the extent and the full range of triggers controlling the potential for transmitter plasticity in these cells are unknown, we show that there are certain cells populations that reliably turn on VAChT translation (Fig. 4B and C), some in response to aging (Fig. 4D to F). How this is accomplished will require further study; but there are many examples of regulated miR degradation (*25*), one of which has been shown to control miR-190 levels (*26*). It is also interesting to consider whether posttranscriptional processes may provide a more general mode of fast but transient control of transmission. We note that there are high levels of *VGAT* transcription in Glu^ACh^ neurons (Fig. 1D). Transient modulation of co-transmission provides a powerful mechanism for sculpting behavior in response to external and internal signals.

## Supporting information

plasmid map

## Acknowledgments

We thank Ed Dougherty in the Brandeis Imaging Facility for assistance. We also thank Paul Garrity, Piali Sengupta and Sebastian Kadener for critical comments on this manuscript.

## Funding

This work was supported by NIH R01067284, NIH R21NS096414 and NIH P01NS090994 to LCG. MH and EJRR were supported by NIH T32NS019929 and EJRR was supported by NIH F31NS110273. Stocks obtained from the Bloomington Drosophila Stock Center (NIH P40OD018537) were used in this study.

## Author contributions

Conceptualization: LCG, YZ, NC

EJRR Methodology: YZ, NC, EJRR, MH, AY

Investigation: YZ, NC, EJRR, AY

Visualization: YZ, NC, EJRR, AY

Funding acquisition: LCG

Supervision: LCG

Writing – original draft: NC, LCG

Writing – review & editing: NC, YZ, MR, LCG

## Competing interests

The authors declare no competing interests.

## Data and materials availability

The data generated and analyzed during this study are available from the corresponding author on request.

## Supplementary Materials

### Materials and Methods

#### Fly strains and husbandry

All flies were raised on standard food at 25°C with a 12h:12h light-dark cycle, except for the *Tubulin-Gal80^ts^* experiments to induce expression at different developmental stages, where flies were raised at either 18°C or 29°C. Male and female flies were collected at eclosion and aged as specified before performing experiments. *VT030559-GAL4* was obtained from Vienna Drosophila Resource Center (VDRC) stock center. *VAChT^[MI08244]^* (#55439), *nsyb-Gal4* (#51941), *VGluT-Gal4* (#60312), *VGluT-p65-AD* (#82986), *VGluT-GAL4-DBD* (#60313), *ChAT-Gal4* (#60317), *GH146-Gal4* (#30026), *GMR81C04-Gal4* (#48378), *VGAT-Gal4* (#84696), *UAS-miR-190-sponge* (#61397), *UAS-scramble-sponge* (#61501), *UAS-Flp* (#4539), *UAS-CD4-GFP^1-10^* (#93016), *VGluT-Gal80* (#58448) and *tubulin-Gal80^ts^* (#7016) were obtained from Bloomington *Drosophila* stock center. *UAS-myrGFP-2A–RedStinger* (*27*) was obtained from the Ganetzky lab at University of Wisconsin, and *UAS-UNC84::GFP* from Gilbert Henry at Janelia Research Campus.

#### Generation of EGFP::VAChT, RFP::VGluT and RFP::VGAT lines

To knock in *EGFP* at the N-terminal of *VAChT*, we designed a guide RNA which recognized the beginning of *VAChT* with an online tool (http://targetfinder.flycrispr.neuro.brown.edu/) and created a donor plasmid (pMC1-EGFP-VAChT plasmid in Data S1). The guide RNA was cloned into a pU6 plasmid (Addgene, #45946) and injected into Cas9 flies (*y,sc,v; nos-Cas9/CyO; +/+*) with the donor plasmid. By the same strategy, we knocked in *RFP* at the N-terminal of *VGluT* and *VGAT*. All guide RNAs are listed in Table S1 and donor plasmids are shown in Data S1. Correct integrations were confirmed by PCR and sequencing with primers which bind outside the regions of the integrated junction.

#### Creation of Frt-stop-Frt-ECFP::VAChT, Frt-stop-Frt-EGFP::VGluT and Frt-stop-Frt-EGFP::VGAT flies

For the *Frt-stop-Frt-ECFP::VAChT* fly strain, we used the same guide RNA as *EGFP::VAChT* and made a donor plasmid (pMC10-Frt-stop-3p3-RFP-Frt-ECFP::VAChT plasmid in Data S1). We amplified the stop sequence which was flanked by two Frt sites, *ECFP* sequence, and *VAChT* sequence. 3P3 RFP sequence was amplified and inserted between stop and the second Frt site for screening. These fragments were assembled in order and cloned into the pMC10 plasmid. The guide RNA was cloned into pU6 plasmids and injected into Cas9 flies with the donor plasmid. F1 progeny with RFP markers were selected as candidates, and further confirmation was performed by PCR and sequencing. By the same strategy, we made *Frt-stop-Frt-EGFP::VGluT* and *Frt-stop-Frt-EGFP::VGAT* flies. The guide RNAs are listed in Table S1, and the donor plasmids were shown as pMC10-Frt-stop-3P3-RFP-Frt-EGFP::VGluT and pMC10-Frt-stop-3P3-RFP-Frt-EGFP::VGAT in Data S1.

#### Creation of *Flp-VAChT-3’UTR* flies

For the *Flp-VAChT-3’UTR* fly strain, we used the same guide RNA as *EGFP::VAChT* and made a donor plasmid (pMC10-Flp-VAChT-3’UTR plasmid in Data S1). The guide RNA was cloned into the pU6 plasmid and injected into Cas9 flies with the donor plasmid. Correct integrations were confirmed by PCR and sequencing.

#### Creation of split-Gal4 lines

To make the *VAChT-AD* and *VAChT-DBD* fly strains, the phase 0 T2A-p65AD-Hsp70 plasmid (Addgene, #62914) and T2A-Gal4DBD-Hsp70 plasmid (Addgene, #62903) were injected into *VAChT^[MI08244]^* flies with pBS130 plasmid (Addgene, #26290) which encodes phiC31 integrase. Progeny were crossed to *yw* flies to check for spGAL4 insertion. Male flies with yellow marker were selected as candidates, and then checked by PCR to obtain insertion lines in the correct orientation.

For the *VGAT-AD* and *VGAT-DBD* lines, we first made a *3P3-RFP-VGAT* fly strain utilizing the same guide RNA as *RFP::VGAT* (Table S1) and a donor plasmid which contained attp flanked *3P3-RFP* sequences (fig. S1A). Flies were first screened for RFP expression, and then confirmed by PCR and sequencing. To make *VGAT-AD* flies, the AD sequence was amplified from T2A-p65AD-Hsp70 plasmid (Addgene, #62914) and attached at the N terminal of the VGAT sequence. The whole AD sequence which was flanked by two inverted-attB sites was cloned into the pBS-KS-attB2 plasmid (Addgene, #62897). This plasmid was injected into *3P3-RFP-VGAT* flies, with plasmids that expressed phiC31 recombinase. By the same strategy, we made *VGAT-DBD* flies using T2A-Gal4DBD-Hsp70 plasmid (Addgene, #62903). F1 progeny without RFP marker were selected as candidates, and further confirmation by PCR and sequencing were performed.

#### Creation of VAChT-GFP^1-10^, VAChT-GFP^11^, and VGAT-GFP^11^ lines

To make the *VAChT-GFP^1-10^* and *VAChT-GFP^11^* fly strains, we first chosen a luminal-side insertion site using *in silico* prediction (https://phobius.sbc.su.se/). We used the same guide RNA as *EGFP::VAChT*, and created donor plasmids (VAChT-GFP1-10 plasmid and VAChT-GFP11 plasmid in Data S1). The guide RNA was cloned into a pU6 plasmid and injected into Cas9 flies with the donor plasmids. Correct integrations were confirmed by PCR and sequencing.

For the *VGAT-GFP^11^* line, a luminal-side insertion site was chosen using *in silico* prediction (https://phobius.sbc.su.se/). The GFP^11^ sequence was inserted at the last luminal side site of the VGAT. The whole sequence was flanked by two inverted-attB sites, and cloned into the pBS-KS-attB2 plasmid (Addgene, #62897). This plasmid (VGAT-GFP11 plasmid in Data S1) was injected into *3P3-RFP-VGAT* flies showed above, with plasmids that expressed phiC31 recombinase. F1 progeny without RFP marker were selected as candidates, and further confirmation by PCR and sequencing were performed. Luminal location of the tags was confirmed as shown in fig. S11.

#### Creation of UAS-ChAT, UAS-VAChT, UAS-Fluc-ChAT 3’UTR and UAS-Fluc-ChAT del lines

For the *UAS-RFP::ChAT* fly strain, the coding region of ChAT was amplified from a *Canton-S* wild type fly cDNA library, and inserted into the pUAST-attB plasmid (Addgene, 8489bp) using the Gibson assembly method (UAS-RFP::ChAT plasmid in Data S1). To allow visualization of ChAT expression, RFP was inserted in-frame before the ChAT coding region. Using the same strategy, GFP1-10 and VAChT coding regions were amplified and inserted into pUAST-attB to make the *UAS-VAChT* fly line (UAS-GFP1-10::VAChT plasmid in Data S1).

For the *UAS-Fluc-ChAT 3’UTR* fly line, we amplified the Fluc sequence from the Ac/Fluc plasmid (a gift of Ravi Allada) and the ChAT 3’UTR sequence from the *Canton-S* wild type fly genome. These sequences were assembled in order and cloned into the pUAST-attB plasmid (UAS-Fluc-ChAT 3’UTR plasmid in Data S1). For the *UAS-Fluc-ChAT del* fly line, the same sequences were used, except that the predicted miR-190 binding sites were removed from ChAT 3’UTR (UAS-Fluc-ChAT del plasmid in Data S1).

All plasmids were checked by sequencing. UAS-RFP::ChAT, UAS-Fluc-ChAT 3’UTR and UAS-Fluc-ChAT del plasmids were injected into *phiC31-attP* flies (Bloomington Stock Center #79604) which have an attP site on the second chromosome to allow targeted integration. UAS-GFP1-10::VAChT plasmid was injected into *phiC31-attP* flies (Bloomington Stock Center #8622), which have an attP site on the third chromosome. The progeny of injected flies was screened for *w^+^* red eye marker, and then checked by PCR and sequencing.

#### INTACT purification of nuclei

Nuclei from *Glu^ACh^>UNC84::GFP* and *GABA^ACh^>UNC84::GFP* heads were prepared according to the INTACT protocol (*13*), with some adjustments. Briefly, whole flies were flash frozen on dry ice in 15 ml tubes and vortexed for 5 cycles of 15 s vortexing at max speed and 1 min of resting on dry ice. Heads were separated from bodies using frozen No.40 and No.25 brass sieves. Sieved heads were placed in pre-chilled 1 ml dounce homogenizers and homogenized using a modified INTACT lysis buffer (10mM Tris-HCl pH7.5, 2mM MgCl2, 10mM KCl, 0.6mM Spermidine, 0.2mM Spermine, 1mM DTT, 0.03% Tween-20, 1% BSA, 1x cOmplete Protease inhibitor), for 15 strokes with Pestle A and 15 strokes of Pestle B. Homogenized lysate was filtered through a 20 µm CellTrics Filter (Sysmex Flow Cytometry), centrifuged for 5 min at 800 RCF. Supernatant was removed and lysate was resuspended in modified INTACT lysis buffer and filtered through a 10 µm CellTrics Filter (Sysmex Flow Cytometry). Filtered lysate was then subject to anti-GFP immunoprecipitation and RNA extraction as previously described (*13*).

#### RNA-seq and data analysis

Purified RNA was subject to PolyA enrichment using the Poly(A)Purist Mag Kit (Thermofisher) according to protocol. Purified Poly(A) RNA was quantified using the Qubit 2.0 RNA HS Assay (Thermofisher), and 10 ng of RNA per sample was used for library prep using the NextFlex Rapid Directional qRNA-Seq Kit 2.0 (PerkinElmer) and sequenced on a NextSeq 550 using the 75 cycle High Output Kit (Illumina).

UMIs were extracted and appended to reads from sequenced libraries using umI_tools extract with the following parameters: --bc-pattern=NNNNNNNNN --bc-pattern2=NNNNNNNNN. Processed reads were then aligned against the dm6 reference genome with STAR using the following parameters: --outFilterMismatchNoverLmax 0.05 --outFilterMatchNmin 15 – outFilterMultimap Nmax 1 --outSJfilterReads Unique --alignMatesGapMax 25000. Aligned reads were converted to BAM files and sorted using samtools, and were deduplicated using umi_tools dedup. Reads were counted using featurecounts, and normalization and differential expression was conducted using Deseq2. The full data set is available at NCBI; GEO accession number GSE221859

#### Immunohistochemistry and image processing

For dissection and staining of adult fly brains, the protocol from Janelia (https://www.janelia.org/project-team/flylight/protocols) was used. Briefly, brains were dissected in S2 solution, and then fixed in 2% PFA solution for 55 min at room temperature (RT). Then the samples were washed 4×10 mins by 0.5% PBST solution, and blocked with 5% goat serum in PBST solution for 1.5 hours. After that, the samples were incubated in primary antibody solutions for 4 hours at RT and continued incubation at 4°C for over two nights. Then samples were washed 3×30 min by 0.5%PBST, incubated in secondary antibody solutions for 4 hours at RT, with continued incubation at 4°C for over three nights. The same washing protocol was performed after secondary antibody incubation, then fixed by 4% PFA again for 4h at RT and mounted in Vectashield mounting medium (Vector Laboratories).

The primary antibodies used were: rabbit anti-RFP (1:200, Takara), rabbit anti-GFP (1:1000, Thermo Fisher), mouse anti-GFP (1:200, Sigma), mouse anti-Brp (1:100, DSHB), anti-VGluT(*28*) (1:200, generous gift from Aaron DiAntonio, Washington University) and anti-VGAT (*29*) (1:200, generous gift from David Krantz, UCLA). Alexa Fluor 488 anti-mouse/rabbit antibody (Invitrogen) and Alexa Fluor 635 anti-mouse/rabbit antibody (Invitrogen) were used as secondary antibodies at 1:200 dilutions. All images were taken using Leica SP5 confocal microscope under 20x or 60x objective lens. Then the pictures are processed and analyzed using ImageJ Fiji software(*30*).

#### Sleep and locomotor activity

Individual 3-5 day old male flies were loaded into 65mm x 5mm glass tubes (Trikinetics, Waltham, MA) using CO_2_ anesthesia. One end of the tube is food containing 5% agarose and 2% sucrose, the other side is a cotton ball to cover it. The flies were entrained under standard 12:12 light-dark conditions for 2 days prior to data collection.

Locomotor activity was collected with the *Drosophila* Activity Monitoring System (Trikinetics) as previously described (*31*). Sleep is defined as consecutive inactivity for five or more minutes (*32*). All sleep parameters, including total sleep duration, number of sleep episodes and mean episode duration were analyzed using an Matlab program described previously (*31*) and averaged across 4 days. Statistical analysis was performed with GraphPad Prism. For all sleep parameters a D’Agostino & Pearson test was used to determine normality of data. If data were normally distributed they were analyzed using a Student T-test or ANOVA followed by Tukey test for multiple comparisons (depending on the number of groups). If data were not normally distributed they were analyzed using a Mann-Whitney or Kruskal-Wallis test followed by Dunn’s test for multiple comparisons.

#### S2 cell assay (*33*)

S2 cells in 12-well plates were cotransfected with 15 ng of Ac/Fluc (or its derivatives), 15 ng of Ac/Rluc, and 270 ng of Ac/miR-190 or Ac/scramble by Effectene transfection reagent (Qiagen). Ac/Fluc derivatives included Fluc with ChAT-3’UTR, Fluc with VAChT-3’UTR, and Fluc with ChAT-3’UTR with the three predicted miR-190 binding sites removed. The primers are listed in Table S2. Cells were harvested 48 hours after transfection and a dual luciferase assay was performed (Promega).

#### *In vivo* Luciferase assays

15 male fly brains were collected for each sample, then homogenized in 100 µl Promega Glo Lysis Buffer (Promega, Cat# E2510) at room temperature. Homogenized samples were incubated for 10 min at room temperature, and then centrifuged for 5 min to pellet the brain remains. 50 µl of supernatant was transferred to an Eppendorf tube on ice, and another 450 µl lysis buffer was added. A multichannel pipette was used to transfer 20 µl of each sample to a white-walled 96-well plate (Costar), then 20 µl Promega Luciferase Reagent (Promega, Cat# E2510) was added to each well. The plate was incubated in dark for 10 min. Luminescence was measured on a Luminometer plate reader (Promega, Cat# GM3000).

---

**Figure S1.**
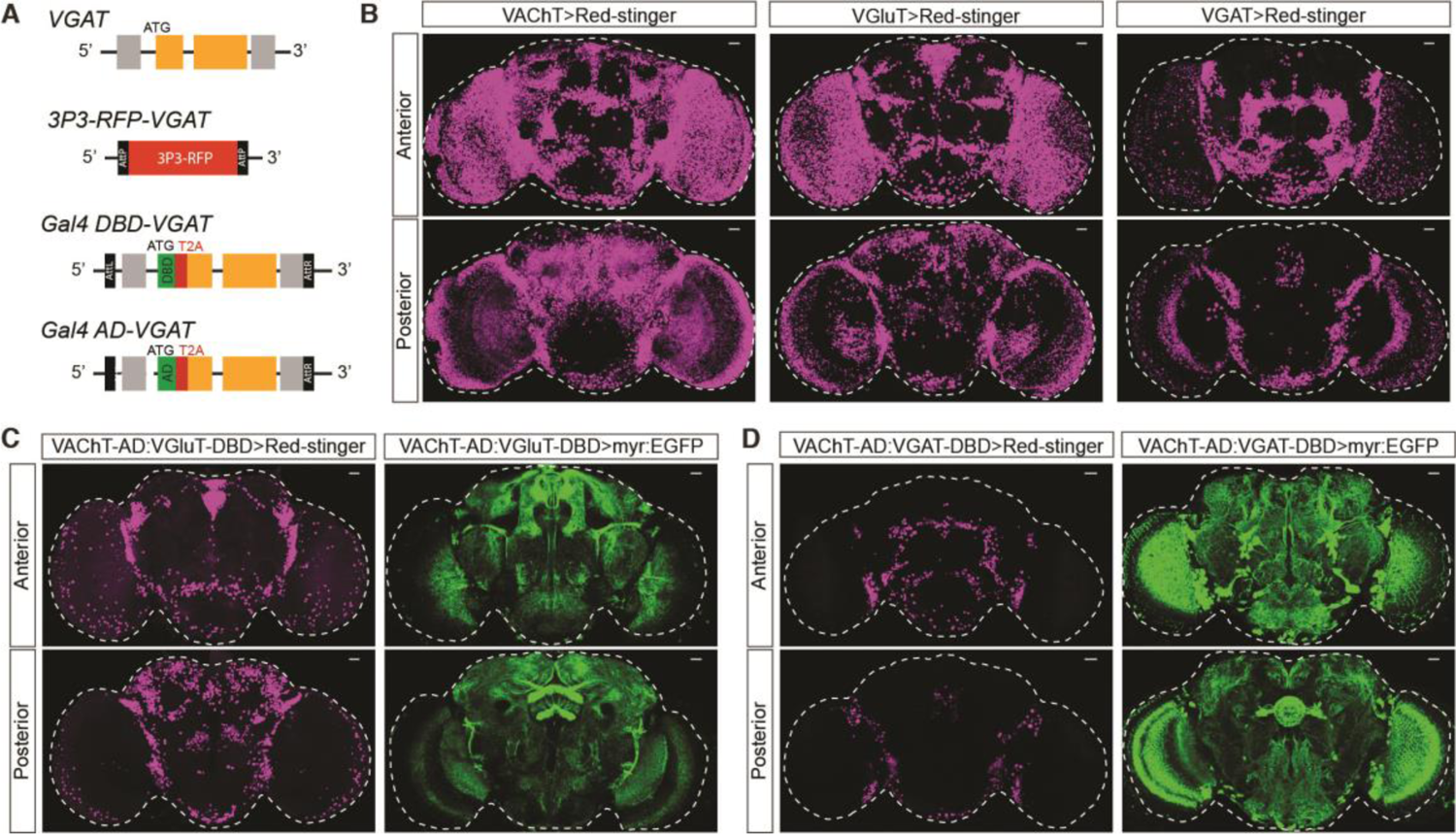
Expression patterns of *VAChT:VGluT* and *VAChT:VGAT* split Gal4s. (**A**) Schematic diagram showing fused *Gal4-DBD* and *Gal4-AD* knock-in strategy: attp-flanked 3P3-RFP was knocked in to replace the whole *VGAT* gene using CRISPR/Cas9. This cassette was then is replaced by attb-DBD-T2A-*VGAT*-attb or attb-AD-T2A-*VGAT*-attb using *phiC31* recombination. Grey bars indicate the UTRs, while yellow bars indicate exons. (**B**) Magenta shows the soma (nuclei) of *VAChT-Gal4, VGluT-Gal4* and *VGAT-Gal4* expression patterns: anterior views (top) and posterior (bottom). (**C-D**) Magenta shows the somatic regions of *VAChT-AD:VGluT-DBD* split-Gal4 (**C**) and *VAChT-AD:VGAT-DBD* split-Gal4 (**D**) flies, while green shows the neuronal projection regions. Dashed white lines indicate the whole brain outline. Scale bars = 20µm for each panel. Comparison of the number of cells in B vs C/D shows that the split-GAL4s represent only a subset of the neurons captured by the broader drivers.

**Figure S2.**
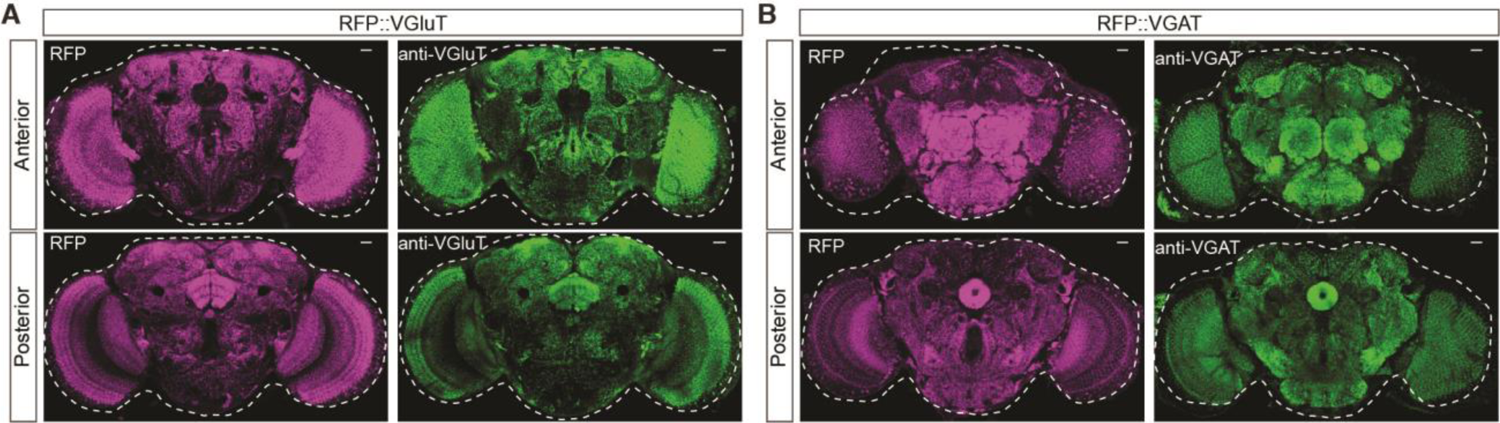
Validation of fusion alleles. To validate the expression patterns of our tagged vNTs, we stained heterozygous animals which have one FP-tagged allele and one untagged allele with anti-VGAT or anti-VGluT. RFP::VGluT protein from our fusion allele overlaps with wildtype chromosome VGluT staining in *RFP::VGluT/+* animals (**A**). RFP::VGAT protein overlaps with wild type chromosome VGAT staining in *RFP::VGAT/+* animals (**B**). Anterior (top) and posterior (down) pictures are shown separately. Scale bars = 20µm for each panel.

**Figure S3.**
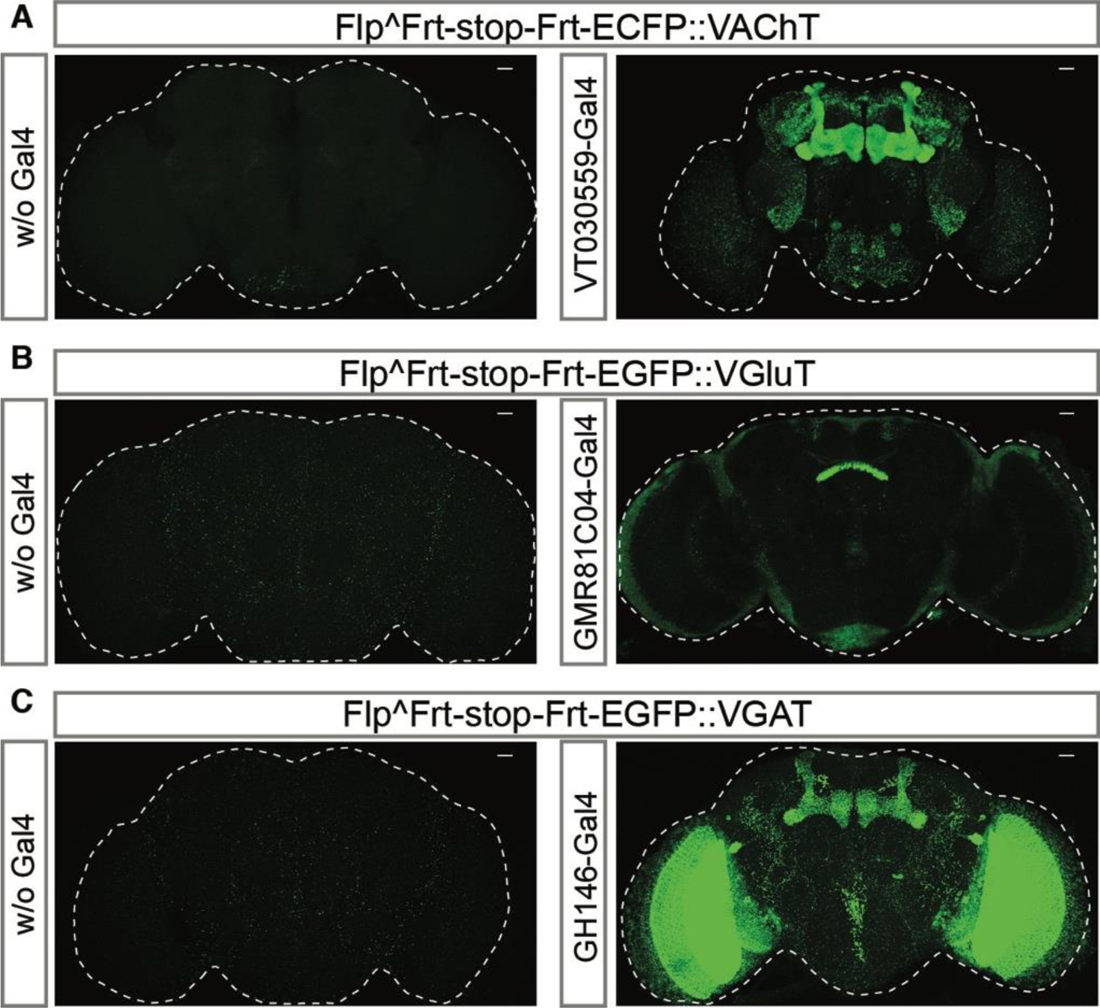
Validation of conditional vNT::FP fusion alleles. Gal-4 drivers for brain regions known to contain neurons expressing a particular neurotransmitter system were used to validate our flip-out strategy. (**A**) *VT030559-Gal4* driving FLP recombinase allows expression of ECFP::VAChT in the mushroom body Kenyon cells, which are known to be cholinergic. No ECFP signal is present without GAL4 expression. (**B**) *GMR81C04-Gal4* driving Flp recombinase allows EGFP::VGluT protein expression in FSB neurons, which are glutamatergic. No EGFP signal is detected when no GAL4 is expressed. (**C**) *GH146-GAL4* driving Flp recombination derepresses EGFP::VGAT protein expression in the APL neurons which are known to be GABAergic. No EGFP signal is detected without GAL4 expression. Dashed white lines indicate the brain outline. Scale bars = 20µm.

**Figure S4.**
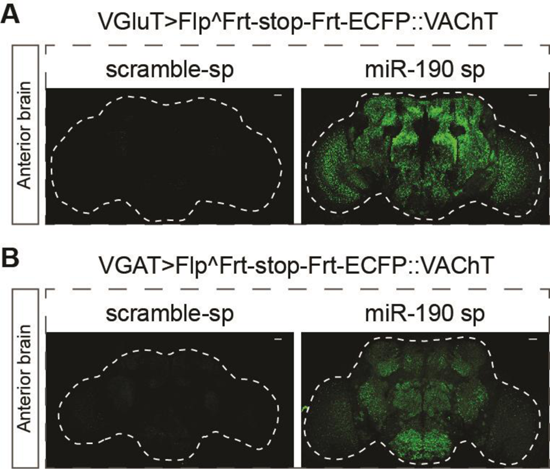
Suppression of miR-190 function allows VAChT protein expression in VGluT (A) and VGAT (B) positive neurons. Representative pictures show the anterior brain signals. Posterior brain stacks are shown in Fig. 2C-D. Dashed white lines indicate the outline of the brain. Scale bars = 20µm.

**Figure S5.**
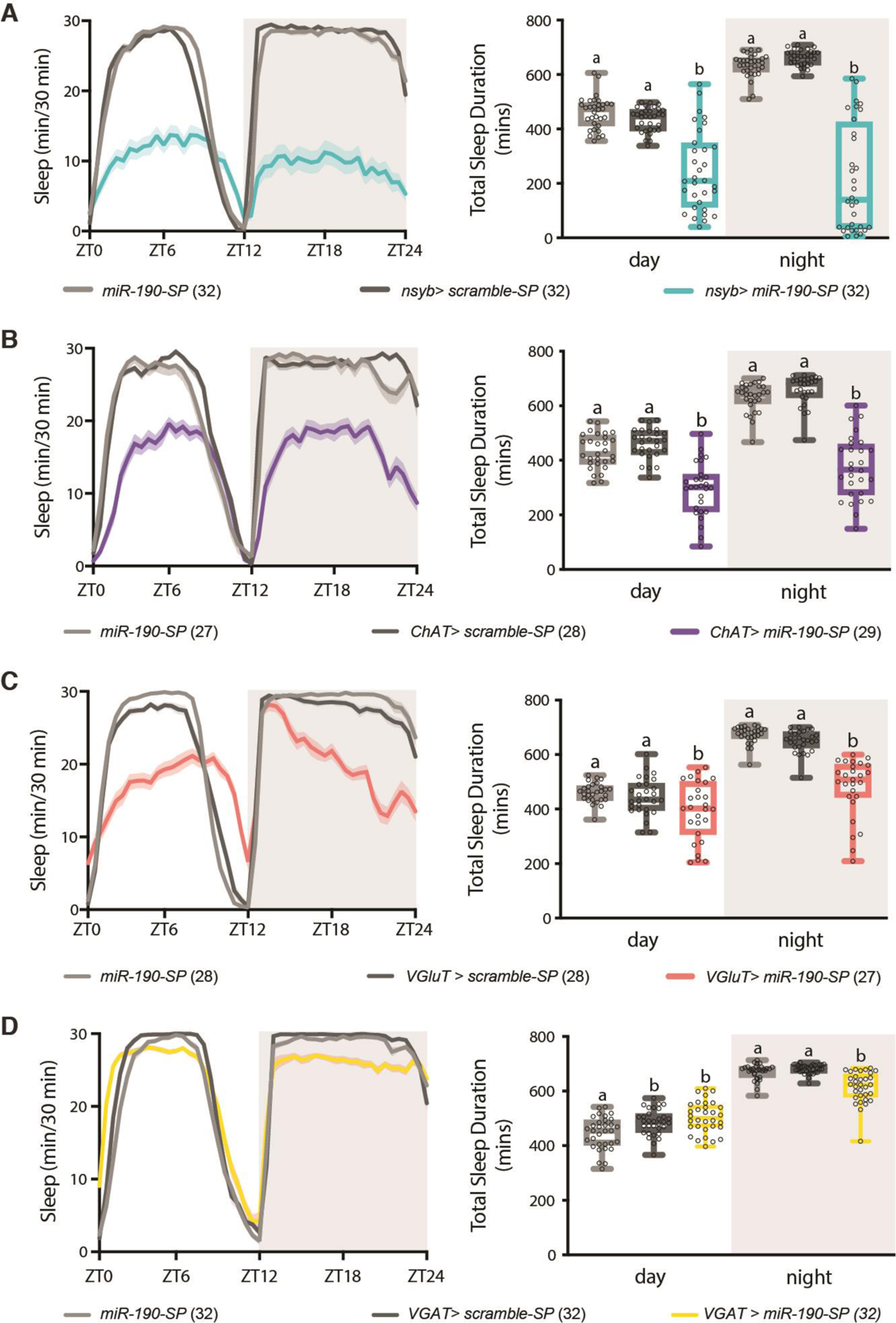
Suppression of miR-190 function reduces total sleep. (**A-D**) Left panels: sleep per 30 mins across 24 hours of a 12:12 light:dark cycle. Right panels: Quantification of total sleep duration when miR-190 function is suppressed in all neurons with *nsyb-Gal4*, a panneuronal driver (**A**), cholinergic neurons with *ChAT-Gal4* (**B**), glutamatergic neurons with *VGluT-Gal4* (**C**), and GABAergic neurons with *VGAT-Gal4* (**D**). Data are shown as mean ± SEM, and gray circles show individual values. Statistical differences are indicated by letters, with genotypes that are not significantly different having the same letter. Data were analyzed with one-way ANOVA with Tukey’s multiple comparisons test or Kruskal-Wallis with Dunn’s multiple comparisons test (depending on data set structure), p < 0.05.

**Figure S6.**
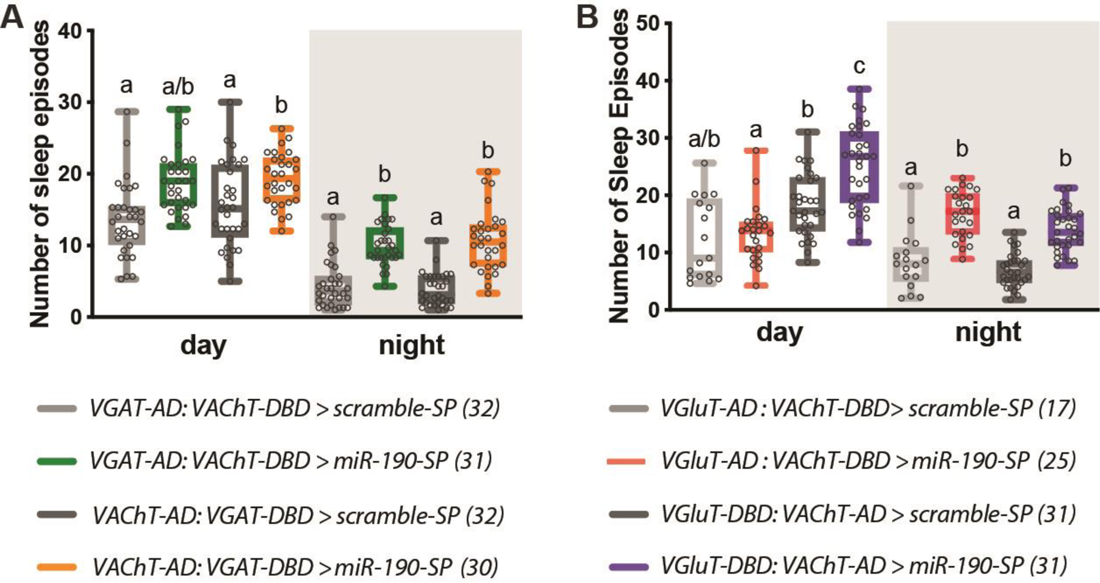
Suppression of miR-190 in GABA^ACh^ and Glu^ACh^ neurons with two independent split-Gal4s fragments sleep and increases the number of sleep episodes. Sleep fragmentation is characterized by both reduced sleep episode duration (as shown in Fig. 3BC) and by increased number of episodes. (**A**) MiR-190 suppression in GABA^ACh^ neurons increases sleep episodes number significantly. (**B**) MiR-190 suppression in Glu^ACh^ neurons makes the number of sleep episodes increase significantly during nighttime. Data are shown as mean ± SEM, and gray circles show individual values. Statistical differences are indicated by letters, with genotypes that are not significantly different having the same letter. Data were analyzed with one-way ANOVA with Tukey’s multiple comparisons test or Kruskal-Wallis with Dunn’s multiple comparisons test (depending on data set structure), p < 0.05.

**Figure S7.**
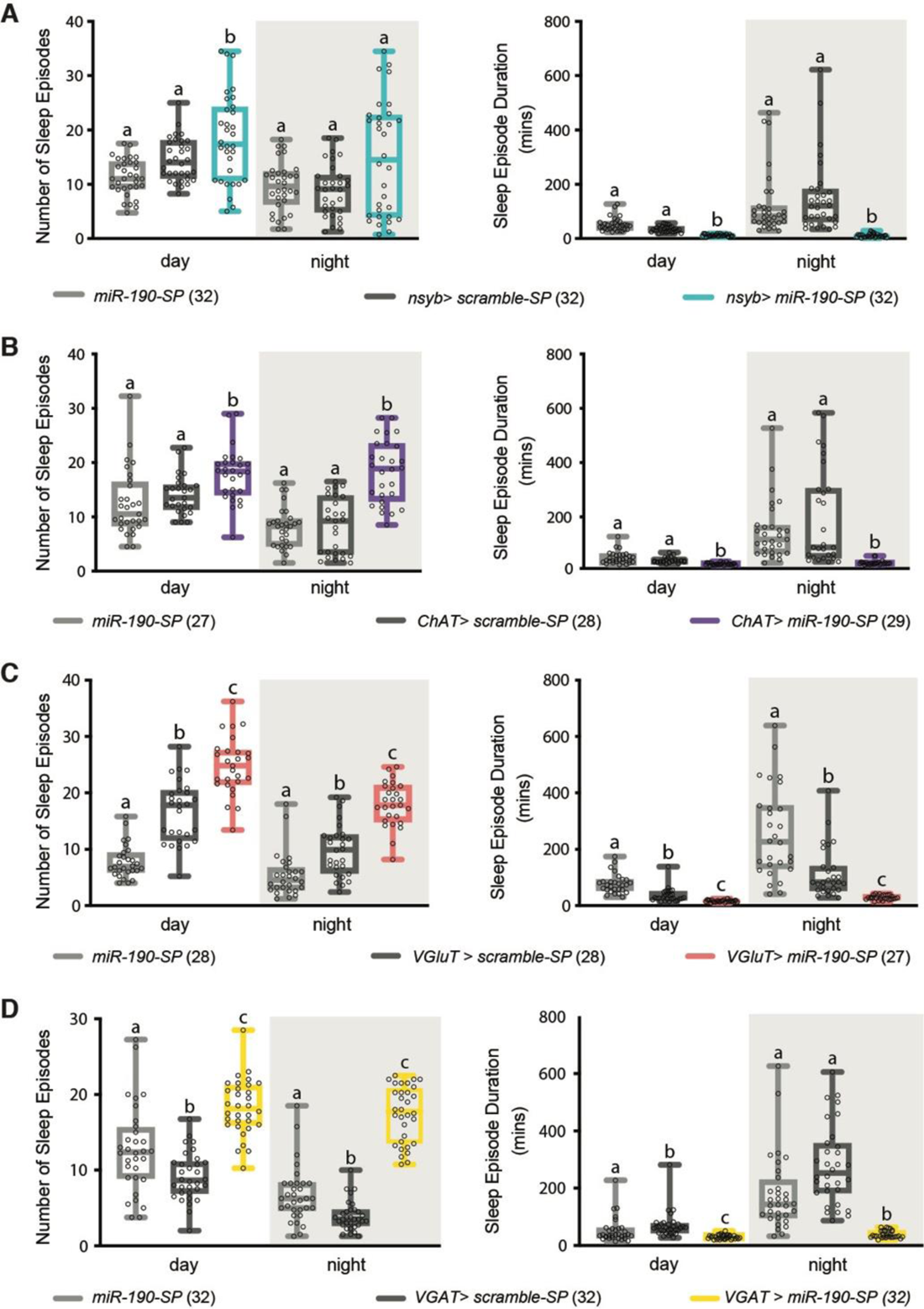
Sleep fragmentation when miR-190 function is suppressed. Quantification of number of sleep episodes (left) and episode duration (right) when miR-190 function is suppressed in all neurons with *nsyb-Gal4* a panneuronal driver (**A**), cholinergic neurons with *ChAT-Gal4* (**B**), glutamatergic neurons with *VGluT-Gal4* (**C**), and GABAergic neurons with *VGAT-Gal4* (**D**). Data are shown as mean ± SEM, and gray circles show individual values. Statistical differences are indicated by letters, with genotypes that are not significantly different having the same letter. Data were analyzed with one-way ANOVA with Tukey’s multiple comparisons test or Kruskal-Wallis with Dunn’s multiple comparisons test (depending on data set structure), p < 0.05.

**Figure S8.**
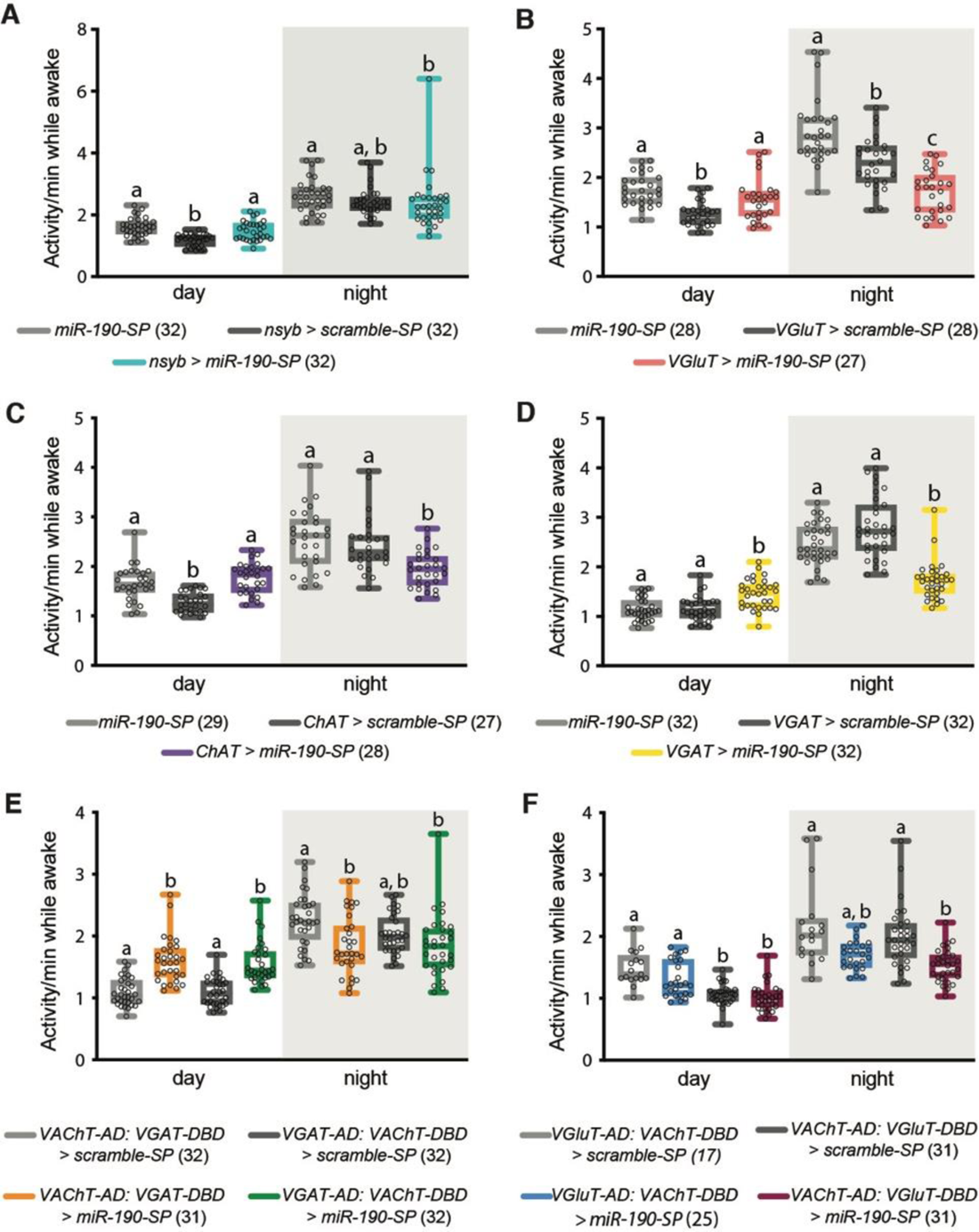
Activity while awake is either not affected or reduced when miR-190 function is suppressed. Quantification of activity while awake when miR-190 function is suppressed in all neurons with *nsyb-Gal4* a panneuronal driver (**A**), glutamatergic neurons with *VGluT-Gal4* (**B**), cholinergic neurons with *ChAT-Gal4* (**C**), GABAergic neurons with *VGAT-Gal4* (**D**), GABA^ACh^ neurons with two different split-Gal4s (**E**), and Glu^ACh^ neurons with two different split-Gal4 drivers (**F**). Data are shown as mean ± SEM, and gray circles show individual values. Statistical differences are indicated by letters, with genotypes that are not significantly different having the same letter. Data were analyzed with one-way ANOVA with Tukey’s multiple comparisons test or Kruskal-Wallis with Dunn’s multiple comparisons test (depending on data set structure), p < 0.05.

**Figure S9.**
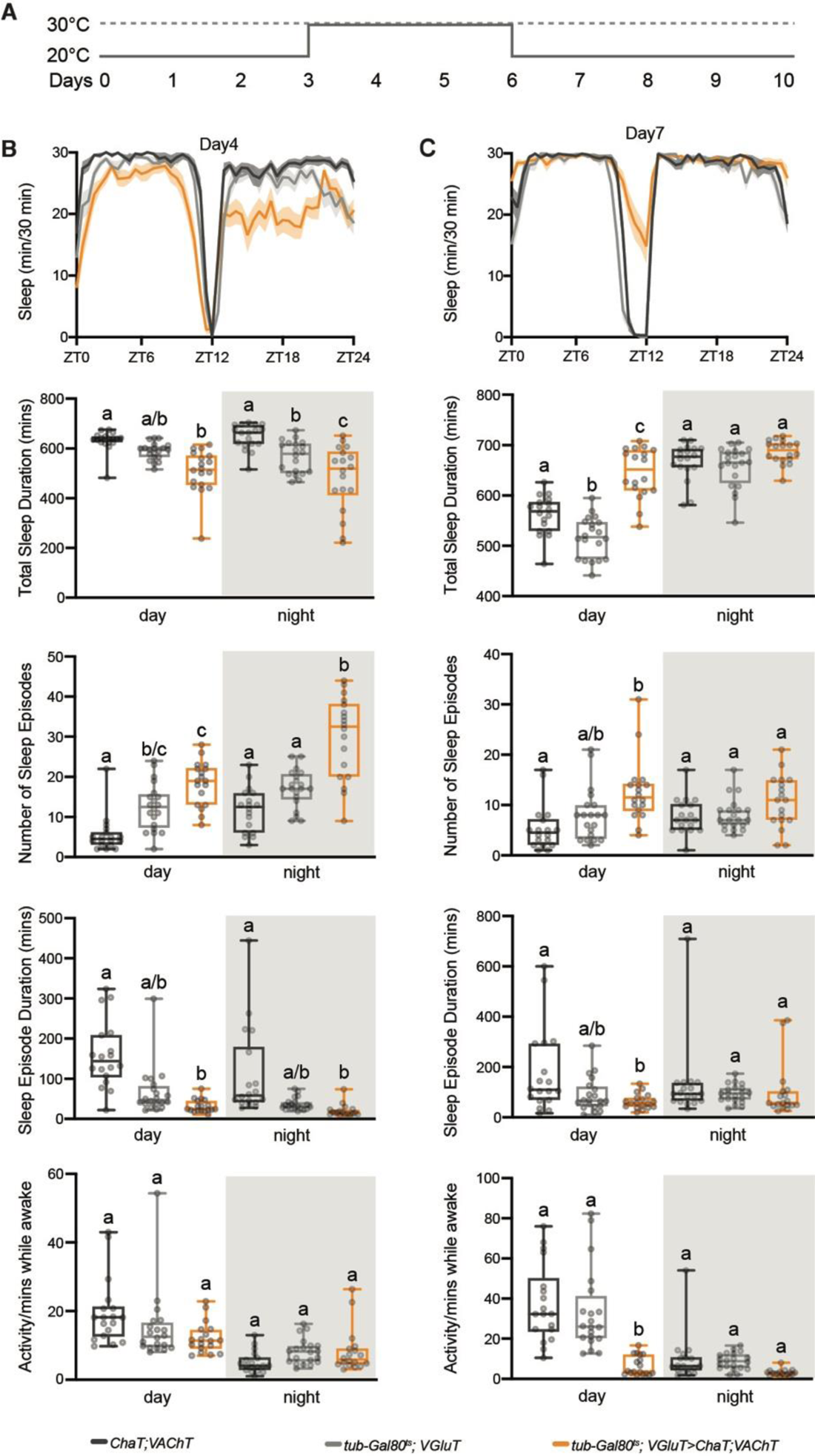
Overexpression of ChAT and VAChT in adult glutamatergic neurons decreases and fragments sleep. (**A**) Schematic diagram of temperature shift to 30°C on day 4 and back to 20°C on day 7. (**B**) Overexpression of ChAT and VAChT in glutamatergic neurons on day 4 decreases nighttime sleep significantly and increases the number of nighttime sleep episodes significantly. (**C**) On day 7, daytime sleep rebounds significantly, overshooting basal levels, though it is notable that there is a suppression of locomotor activity as well. Sleep structure returns to normal. N=18-20. Data are shown as mean ± SEM, and gray circles show individual values. Statistical differences are indicated by letters, with genotypes that are not significantly different having the same letter. Data were analyzed with one-way ANOVA with Tukey’s multiple comparisons test or Kruskal-Wallis with Dunn’s multiple comparisons test (depending on data set structure), p < 0.05.

**Figure S10.**
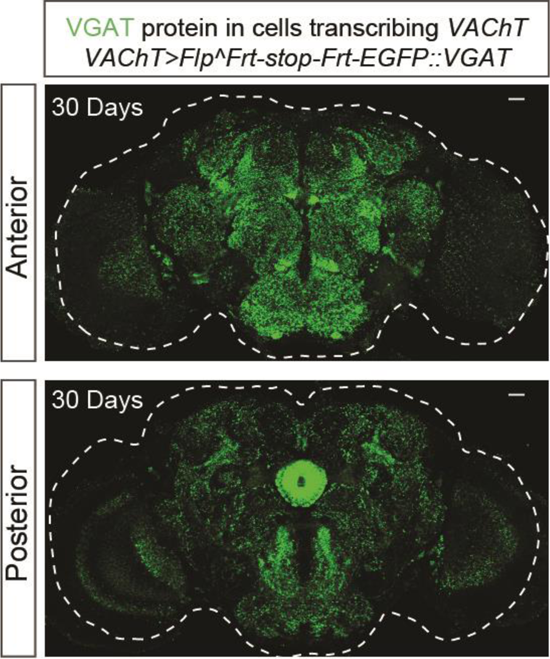
In 30 days aging flies, *VAChT-Gal4* flip-out derepression of *EGFP::VGAT* shows a pattern unchanged by age. Anterior pictures are shown in upper panel, with posterior pictures in lower panel. Young flies are shown for comparison in Fig. 1I. Scale bars = 20µm.

**Figure S11.**
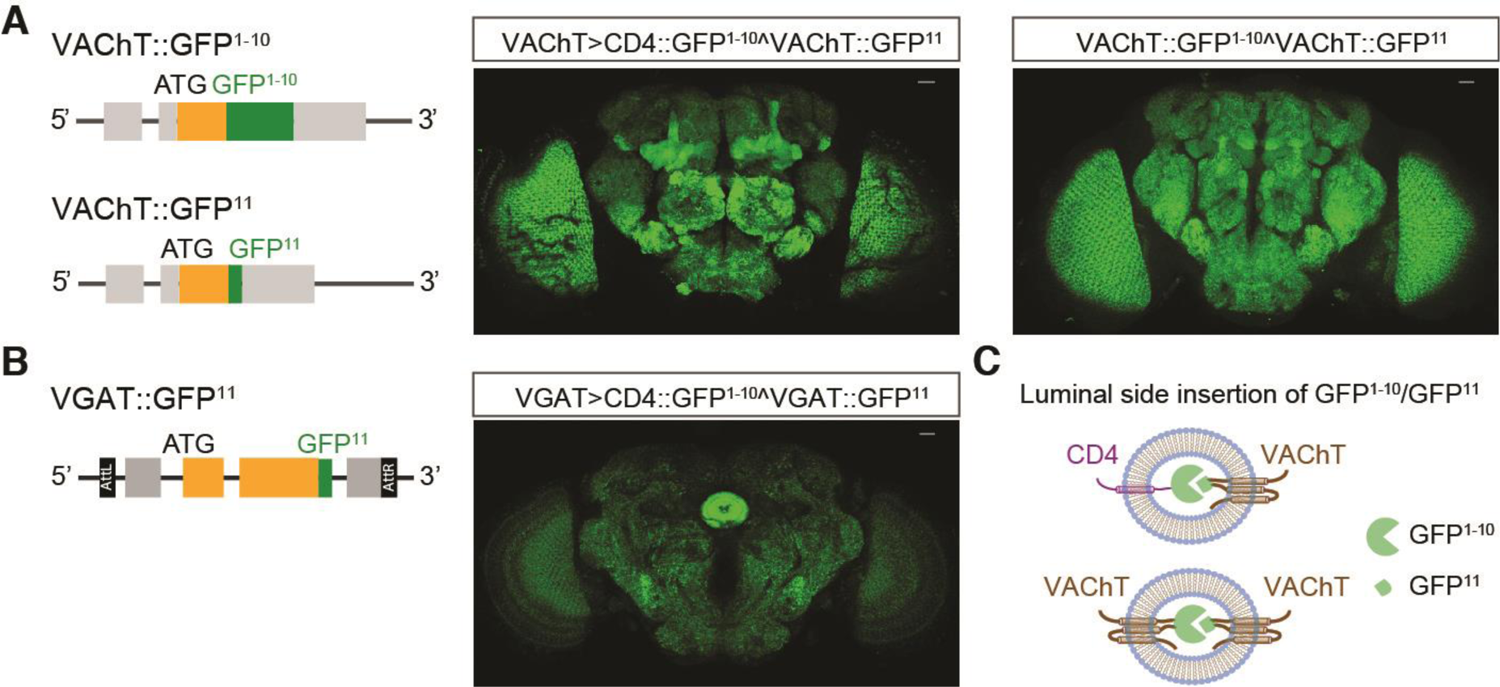
*VAChT::GFP^1-10^*, VAChT::GFP^11^ and VAChT::GFP^11^ display split Gal4 on the luminal side of synaptic vesicles. (**A**) Schematic diagrams in left panel show the *VAChT* insertion sites for *GFP^1-10^* and *GFP^11^* used to create CRISPR alleles. These are predicted to be lumenal (https://phobius.sbc.su.se/). To test this, we crossed *VAChT::GFP^11^* with *VAChT>UAS-CD4-GFP^1-10^* which is known to be lumenal in vesicles. Strong GFP signal confirms GFP^11^ is located on the luminal side. Crossing *VAChT::GFP^1-10^* and *VAChT::GFP^11^* also reconstitutes a strong signal confirming that *VAChT::GFP^1-10^* also displays on the luminal side. (**B**) Schematic diagram in left panel shows the *VGAT* insertion site for *GFP^11^*. Reconstituted GFP signal for *VGAT>CD4-GFP^1-10^* and *VGAT::GFP^11^* indicate the VGAT-GFP^11^ displays on the luminal side. (**C**) The cartoon shows the strategy: only when GFP^1-10^ and GFP^11^ are located on the same side of the vesicle membrane can GFP signal be reconstituted and detected. Scale bars = 20µm.

**Table S1.**
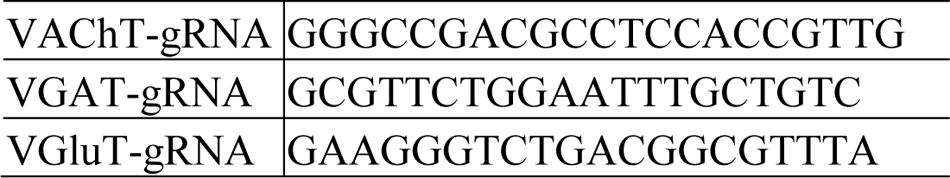
Guide RNAs for VAChT, VGAT and VGluT lines.

**Table S2.**
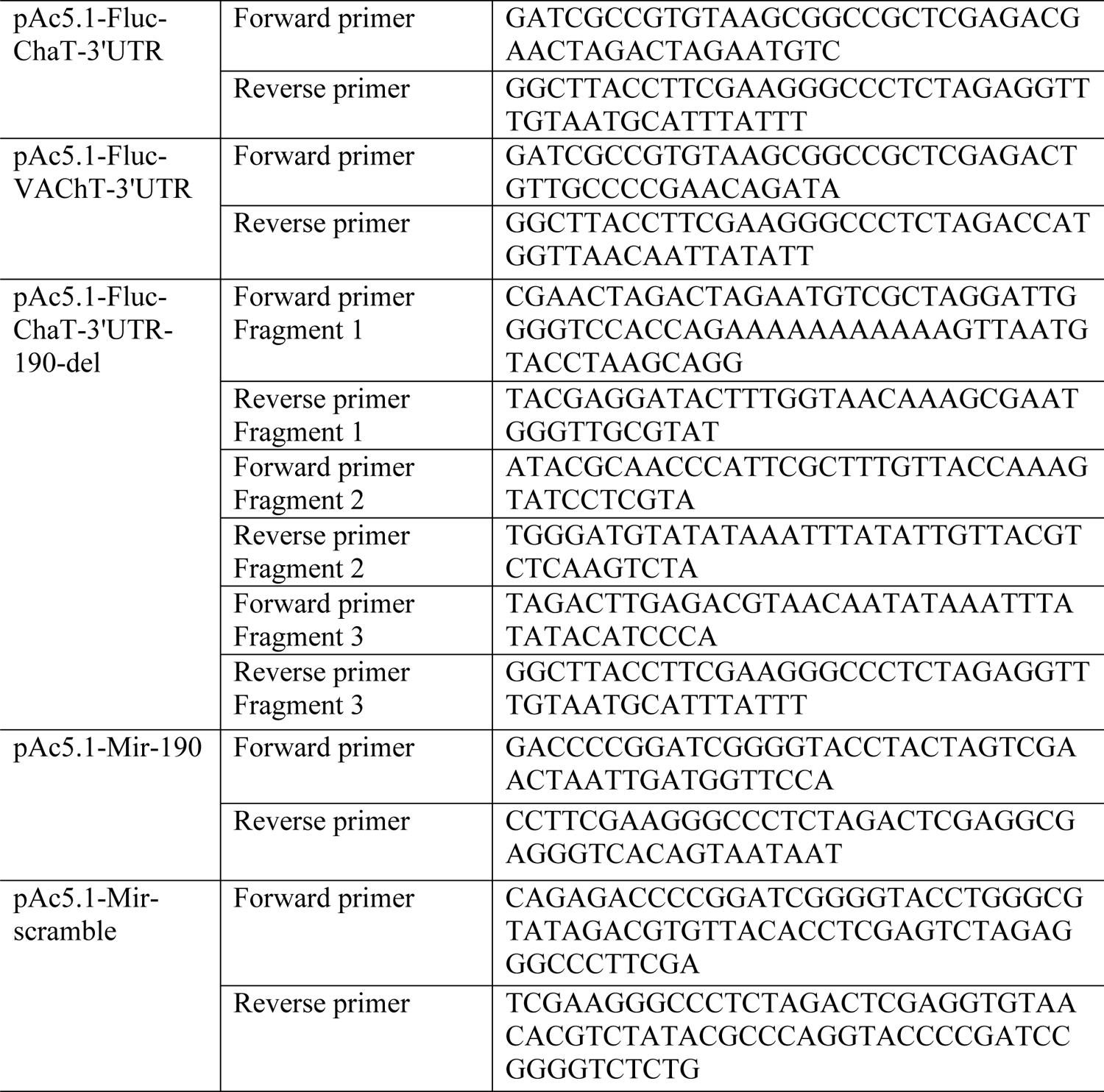
Primers for S2 cell assay plasmids.

**Table S3.**
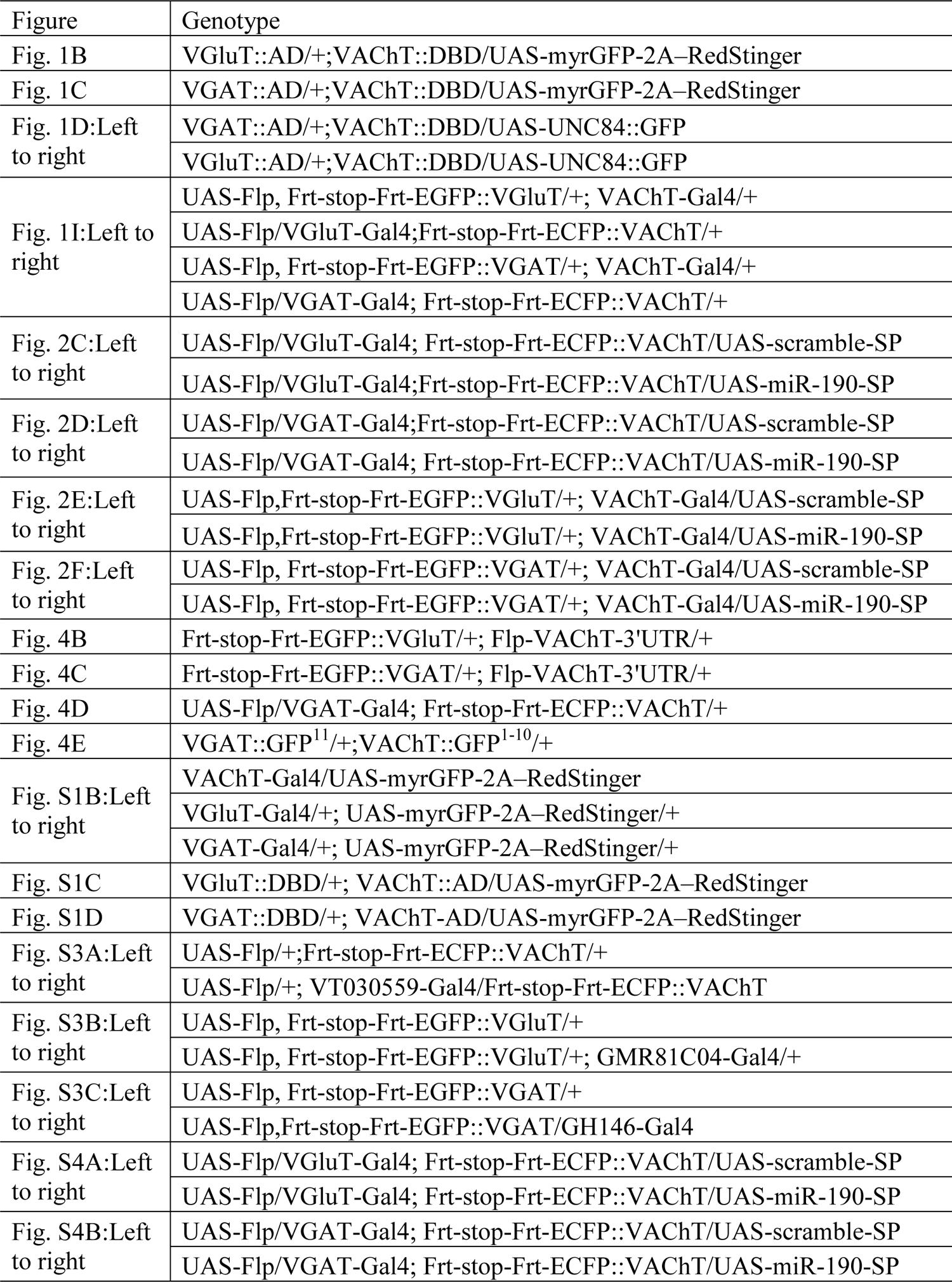
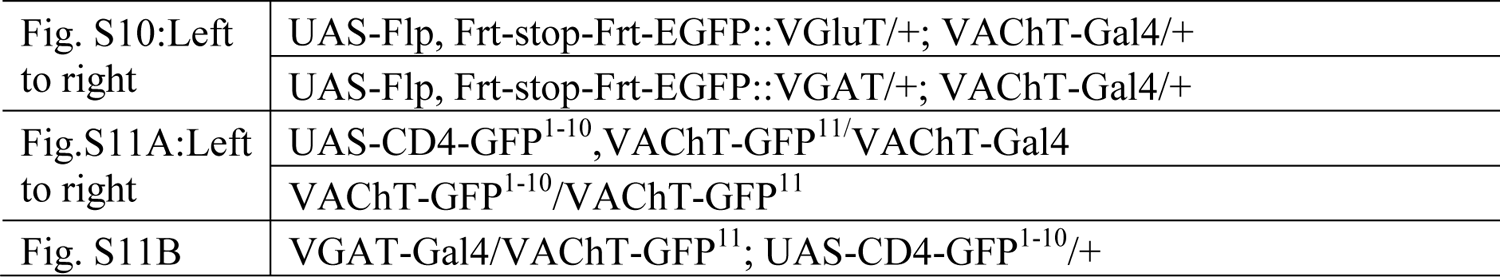
Fly genotypes for figures.

## Data S1

Plasmids maps.

## Notes

### Competing Interest Statement

The authors have declared no competing interest.

## References and Notes

1. N. Ayala-Lopez, S. W. Watts, Physiology and Pharmacology of Neurotransmitter Transporters. Compr Physiol 11, 2279–2295 (2021).

2. K. Silm et al., Synaptic Vesicle Recycling Pathway Determines Neurotransmitter Content and Release Properties. Neuron 102, 786–800 e785 (2019).

3. L. M. Sherer et al., Octopamine neuron dependent aggression requires dVGLUT from dual-transmitting neurons. PLoS Genet 16, e1008609 (2020).

4. N. C. Spitzer, Neurotransmitter Switching in the Developing and Adult Brain. Annu Rev Neurosci 40, 1–19 (2017).

5. A. J. Granger, M. L. Wallace, B. L. Sabatini, Multi-transmitter neurons in the mammalian central nervous system. Curr Opin Neurobiol 45, 85–91 (2017).

6. S. Kim, M. L. Wallace, M. El-Rifai, A. R. Knudsen, B. L. Sabatini, Co-packaging of opposing neurotransmitters in individual synaptic vesicles in the central nervous system. Neuron 110, 1371–1384 e1377 (2022).

7. K. Pankova, A. Borst, RNA-Seq Transcriptome Analysis of Direction-Selective T4/T5 Neurons in Drosophila. PLoS ONE 11, e0163986 (2016).

8. V. Croset, C. D. Treiber, S. Waddell, Cellular diversity in the Drosophila midbrain revealed by single-cell transcriptomics. eLife 7, (2018).

9. H. Lacin et al., Neurotransmitter identity is acquired in a lineage-restricted manner in the Drosophila CNS. eLife 8, (2019).

10. M. P. Nusbaum, D. M. Blitz, E. Marder, Functional consequences of neuropeptide and small-molecule co-transmission. Nat Rev Neurosci 18, 389–403 (2017).

11. A. J. Granger, N. Mulder, A. Saunders, B. L. Sabatini, Cotransmission of acetylcholine and GABA. Neuropharmacology 100, 40–46 (2016).

12. H. Luan, N. C. Peabody, C. R. Vinson, B. H. White, Refined spatial manipulation of neuronal function by combinatorial restriction of transgene expression. Neuron 52, 425–436 (2006).

13. J. Ma, V. M. Weake, Affinity-based isolation of tagged nuclei from Drosophila tissues for gene expression analysis. J Vis Exp, (2014).

14. B. W. Solnestam et al., Comparison of total and cytoplasmic mRNA reveals global regulation by nuclear retention and miRNAs. BMC Genomics 13, 574 (2012).

15. C. Mayr, Regulation by 3’-Untranslated Regions. Annu Rev Genet, (2017).

16. S. Jonas, E. Izaurralde, Towards a molecular understanding of microRNA-mediated gene silencing. Nat Rev Genet 16, 421–433 (2015).

17. T. A. Fulga et al., A transgenic resource for conditional competitive inhibition of conserved Drosophila microRNAs. Nature communications 6, 7279 (2015).

18. G. Artiushin, A. Sehgal, The Drosophila circuitry of sleep-wake regulation. Curr Opin Neurobiol 44, 243–250 (2017).

19. K. Koh, J. M. Evans, J. C. Hendricks, A. Sehgal, A Drosophila model for age-associated changes in sleep:wake cycles. Proc Natl Acad Sci U S A 103, 13843–13847 (2006).

20. E. J. Furshpan, P. R. MacLeish, P. H. O’Lague, D. D. Potter, Chemical transmission between rat sympathetic neurons and cardiac myocytes developing in microcultures: evidence for cholinergic, adrenergic, and dual-function neurons. Proc Natl Acad Sci U S A 73, 4225–4229 (1976).

21. D. Dulcis, P. Jamshidi, S. Leutgeb, N. C. Spitzer, Neurotransmitter switching in the adult brain regulates behavior. Science 340, 449–453 (2013).

22. D. Dulcis et al., Neurotransmitter Switching Regulated by miRNAs Controls Changes in Social Preference. Neuron 95, 1319–1333 e1315 (2017).

23. D. Dulcis, N. C. Spitzer, Reserve pool neuron transmitter respecification: Novel neuroplasticity. Developmental neurobiology 72, 465–474 (2012).

24. M. Bertuzzi, W. Chang, K. Ampatzis, Adult spinal motoneurons change their neurotransmitter phenotype to control locomotion. Proc Natl Acad Sci U S A 115, E9926–E9933 (2018).

25. X. Fu, A. Shah, J. M. Baraban, Rapid reversal of translational silencing: Emerging role of microRNA degradation pathways in neuronal plasticity. Neurobiol Learn Mem 133, 225–232 (2016).

26. C. Y. Shi et al., The ZSWIM8 ubiquitin ligase mediates target-directed microRNA degradation. Science 370, (2020).

27. R. W. Daniels, A. J. Rossano, G. T. Macleod, B. Ganetzky, Expression of multiple transgenes from a single construct using viral 2A peptides in Drosophila. PLoS ONE 9, e100637 (2014).

28. R. W. Daniels et al., A single vesicular glutamate transporter is sufficient to fill a synaptic vesicle. Neuron 49, 11–16 (2006).

29. H. Fei et al., Mutation of the Drosophila vesicular GABA transporter disrupts visual figure detection. J Exp Biol 213, 1717–1730 (2010).

30. J. Schindelin et al., Fiji: an open-source platform for biological-image analysis. Nature methods 9, 676–682 (2012).

31. N. C. Donelson et al., High-resolution positional tracking for long-term analysis of Drosophila sleep and locomotion using the “tracker” program. PLoS ONE 7, e37250 (2012).

32. H. J.C., et al., Rest in Drosophila is a Sleep-like State. Neuron 25, 129–138 (2000).

33. C. Lim et al., The novel gene twenty-four defines a critical translational step in the Drosophila clock. Nature 470, 399–403 (2011).

